# Heritable variation in thermoregulation is associated with reproductive success in the world’s largest bird

**DOI:** 10.1101/2022.03.08.483498

**Authors:** Erik I. Svensson, Mads F. Schou, Julian Melgar, John Waller, Anel Engelbrecht, Zanell Brand, Schalk Cloete, Charlie K. Cornwallis

## Abstract

Organisms inhabiting extreme thermal environments, such as desert birds, have evolved spectacular adaptations to thermoregulate during hot and cold conditions. However, our knowledge of selection for thermoregulation and the potential for evolutionary responses is limited, particularly for large organisms experiencing extreme temperature fluctuations. Here we use thermal imaging to quantify selection and genetic variation in thermoregulation in ostriches (*Struthio camelus*), the world’s largest bird species that is experiencing increasingly volatile temperatures. We found that females that are better at regulating their head temperatures (‘thermoregulatory capacity’) had higher egg-laying rates under hotter conditions. Thermoregulatory capacity was both heritable and showed signatures of local adaptation: females originating from more unpredictable climates were better at regulating their head temperatures in response to temperature fluctuations. Together these results reveal that past and present evolutionary processes have shaped genetic variation in thermoregulatory capacity, which appears to protect critical organs, such as the brain, from extreme temperatures during reproduction.

**Impact Summary:** Large animals inhabiting extreme thermal environments, such as deserts, are predicted to be particularly vulnerable to the increasing temperature fluctuations expected in the future. However, previous work on the evolutionary potential of thermoregulation has primarily focused on the effect of hot temperatures on the survival of small ectotherms. We know little about how large endothermic vertebrates, such as birds, will respond to changing temperatures. Here we study the ostrich (*Struthio camelus*), the world’s largest bird, that inhabits some of the hottest and driest regions on Earth. We show that the ability of females to reproduce during hot conditions is associated with the regulation of their head temperatures (‘thermoregulatory capacity’). Furthermore, variation in thermoregulation is heritable and related to past climatic conditions: females originating from parts of Africa with more extreme temperature fluctuations were better able to thermoregulate, indicating local adaptation to different climatic conditions. Together, these results suggest that thermoregulation in this large desert bird has evolved in response to past climatic conditions, remains genetically variable and is currently under selection through its effect on reproduction.

## Introduction

Organisms need to manage heat and cold stress to survive and reproduce in variable climates (Angiletta, 2009). The universal challenge of coping with thermal stress (Hurley *et al*., 2018; Sales *et al*., 2018; Walsh *et al*., 2019; Schou *et al*., 2021) has promoted the evolution of adaptations to regulate body temperature. For example, body temperature can be maintained by behavioural thermoregulation mechanisms, such as habitat selection (Bogert, 1949; Huey *et al*., 2003; Angiletta, 2009; Muñoz & Losos, 2018; Muñoz, 2022), or by physiological thermoregulation, such as selective brain cooling (Fuller *et al*., 2003) and sweating (Périard *et al*., 2015). Other thermoregulatory mechanisms involve morphological traits, such as the ears of African Elephants (Weissenböck *et al*., 2010) and the bills (Tattersall *et al*., 2009, 2018; Symonds & Tattersall, 2010) and featherless skin patches of birds (McCafferty *et al*., 2011; Szafrańska *et al*., 2020; Gauchet *et al*., 2022). Such morphological structures can act as so called ‘thermal radiators’ or ’thermal windows” that disperse excess heat in an adaptive fashion (Tattersall *et al*., 2009; Symonds & Tattersall, 2010; Weissenböck *et al*., 2010; Darnell & Munguia, 2011; Eastick *et al*., 2019).

Here, we define thermoregulation as the ability of an organism to maintain body temperatures within a relatively narrow thermal zone (Deutsch *et al*., 2008) favourable for survival or reproduction (Huey *et al*., 2003; Muñoz, 2022). This general definition is applicable to both ecto- and endothermic animals and to both physiological and behavioural mechanisms of thermoregulation (Angiletta, 2009). The mechanisms by which animals thermoregulate and how these mechanisms promote survival have been intensely studied. However, we know much less about the quantitative genetics of thermoregulatory traits and how such traits affect reproductive success, particularly for organisms living in the regions most affected by climate change (Walsh *et al*., 2019), such as deserts (Bourne *et al*., 2020; Schou *et al*., 2021).

The evolutionary potential of populations to adapt to thermal stressors, such as heatwaves and cold snaps, depends on genetic variance in traits involved in thermoregulation (Hurley *et al*., 2018; Sales *et al*., 2018; Bourne *et al*., 2020). Current empirical evidence from insects and reptiles suggests that additive genetic variances and hence evolutionary potential in thermal tolerance traits can be high (Ma *et al*., 2014), but are more often low (Hoffmann *et al*., 2003; Kellermann *et al*., 2009; Logan *et al*., 2018; Castañeda *et al*., 2019). In addition, body temperatures and thermal adaptations are typically strongly phylogenetically conserved (Kellermann *et al*., 2012a, 2012b) and evolve slowly compared to other traits (Moreira *et al*., 2021). The late evolutionary biologist George C. Williams even questioned if endothermic body temperatures could evolve at all, suggesting that there may be evolutionary stasis in body temperatures and thermal adaptations (Williams, 1992). However, it is unclear if genetic variance in thermal traits is genuinely low, setting constraints on evolutionary responses, or whether such variation is just difficult to measure, given the challenges in quantifying variation in thermoregulatory capacity across large numbers of individuals. This is a genuine empirical challenge for research on large vertebrates with small population sizes where it is difficult to obtain large sample sizes. To our knowledge there is a lack of quantitative genetic studies on large endothermic animals, which is unfortunate, as they are likely to be particularly vulnerable to changing temperatures.

Large animal bodies have a higher thermal inertia and slower rate of body temperature change compared to small bodies (Bogert, 1949; Angiletta, 2009). Thermal inertia can help maintain body temperatures during cold conditions, but it can also increase physiological stress under sustained heat, jeopardising survival and reproduction (Hurley *et al*., 2018; Sales *et al*., 2018; Bourne *et al*., 2020; Parratt *et al*., 2021; Schou *et al*., 2021, 2022). Large animals are also predicted to be particularly vulnerable to changing climates because their rate of adaptive evolution is potentially limited by longer generation times and lower population sizes (Cardillo *et al*., 2005; Atwood *et al*., 2020). Understanding how large-bodied endothermic animals evolve to cope with thermal stress in fluctuating thermal environments therefore requires particular attention.

Here we study the evolutionary potential of thermoregulation in the world’s largest bird, the ostrich (*Struthio camelus*) (Fig. 1). Estimating additive genetic variance in thermoregulation, and its relationship to fitness, requires measuring these traits across hundreds of individuals of known pedigree. To do this we studied captive breeding populations of ostriches in the Klein Karoo region of South Africa, where a large number of individuals have been reared in a semi-natural environment over 25-years to produce a large nine-generation pedigree (Schou *et al*., 2021, 2022). To quantify thermoregulation, we used thermal imaging (infrared camera technology) which makes it possible to measure the surface temperature of hundreds of individuals in a non-invasive way (Tattersall & Cadena, 2010; McCafferty *et al*., 2015; Gauchet *et al*., 2022). The temperature profiles of 423 adult females were repeatedly measured an average of six times (n_images_ = 2744). Skin surface temperatures have been shown to be significantly and positively correlated with internal body temperatures in studies on both vertebrates and invertebrates (Tsubaki *et al*., 2010; Andreasson *et al*., 2016; Svensson *et al*., 2020; Szafrańska *et al*., 2020; Gauchet *et al*., 2022). In some species of birds, such as red-footed boobies (*Sula sula*), the correlation between skin surface temperatures and internal body temperatures (T_b_) have been found to be very strong, with body temperatures explaining 76-82 % of the variation (Gauchet *et al*., 2022). Thermal imaging has therefore emerged as a powerful and non-invasive technique that has been used to quantify individual variation in thermal profiles and thermoregulatory capacity across diverse taxa under a variety of natural contexts (Tattersall *et al*., 2009; Svensson & Waller, 2013; Andreasson *et al*., 2016; Moore *et al*., 2019; Svensson *et al*., 2020; Szafrańska *et al*., 2020; Gauchet *et al*., 2022).

**Fig. 1.**
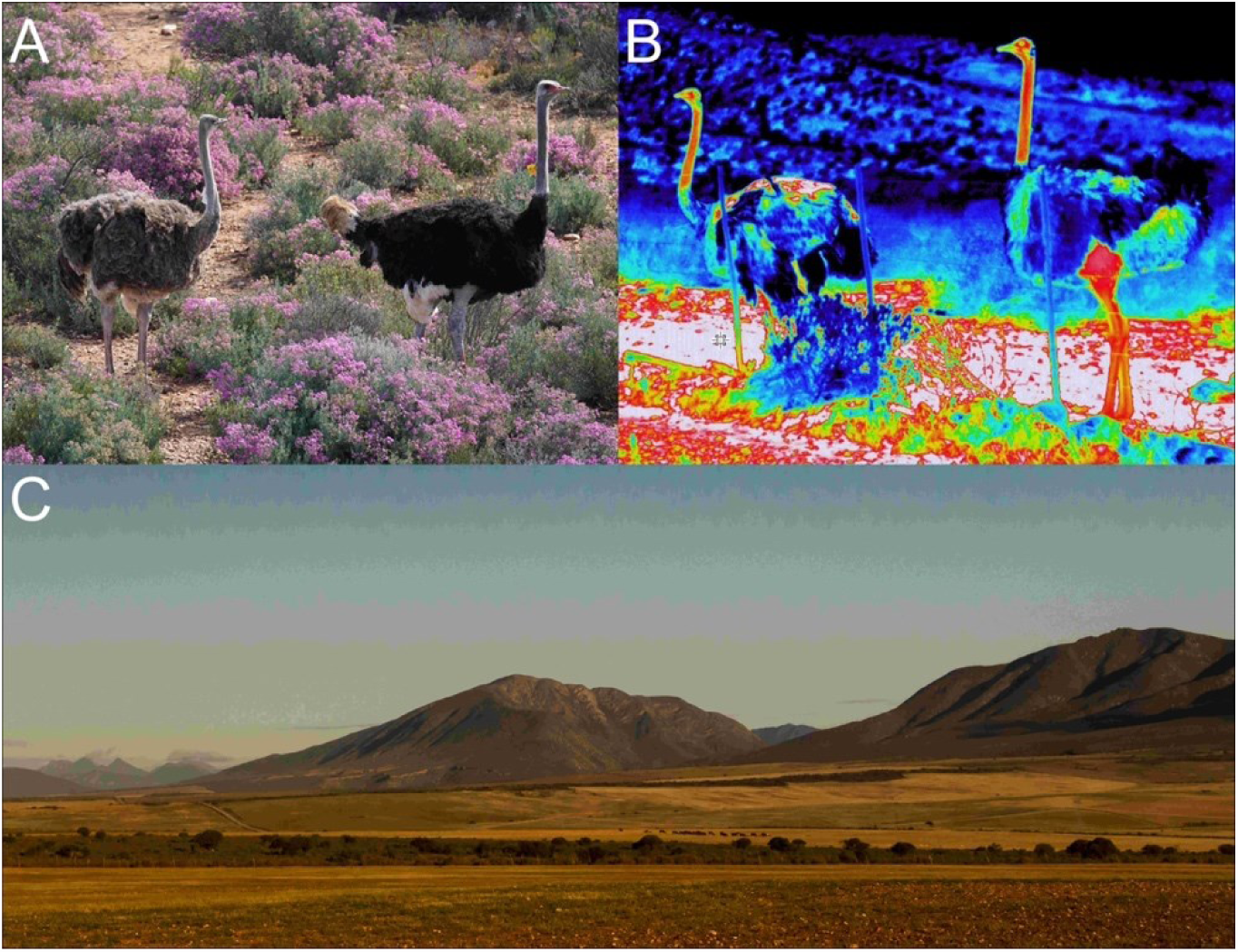
The ostrich inhabits thermally challenging desert environments such as the Karoo in South Africa. **A.** The ostrich (*Struthio camelus*) is the world’s largest living bird with a feathered body and featherless neck and head. *Left*: female. *Right:* male (photograph by C. K. Cornwallis in Karoo National Park, Western Cape Province of South Africa). **B.** Thermal image of a female (left) and male (right) ostrich at our study site in Oudtshoorn (photograph by E. I. Svensson). Note how the long neck is hot and emits excess heat (red, warm colour). **C.** The dry and tree-less semi-desert environment around the study site is characterized by extensive temperature fluctuations ranging from −5°C to 45°C, causing extreme thermal stress (Schou *et al*., 2021) (photograph by E. I. Svensson).

We combined thermal image data with daily measurements of reproductive success and pedigree information to: (1) examine the regulation of head and body surface temperatures across ambient temperatures. We focus specifically on the surface temperature of the head, as shielding the brain from extreme temperatures is of crucial importance for animals (Kilgore *et al*., 1976; Fuller *et al*., 2003; Beltrán *et al*., 2021); (2) test if variation in the efficiency of head thermoregulation influences female reproductive success, which is highly sensitive to temperature (Schou *et al*., 2021, 2022); (3) quantify genetic variation in the efficiency of head thermoregulation; and (4) investigate if ostriches originating from regions with different climatic regimes have evolved differences in their head thermoregulation.

One complication with measuring variation in thermoregulation in free-ranging animals under natural conditions is that it is not possible to control previous exposure to temperature stress, for example, due to variation in microclimatic and biotic factors. A high temperature measurement may therefore be because the animal was exposed to heat, has poor thermoregulatory ability for a given temperature, or both. However, by calculating temperature differences between the head and other body parts it is possible to estimate the ability of individuals to regulate head temperatures for a given thermal stress load (as indicated by the temperature of that body part). A high head temperature *relative to the body* cannot be due to high heat exposure per se as both body parts are exposed to similar external conditions. Consequently, the difference between head and body is likely to capture the specific ability of individuals to regulate their head temperature, a proxy for vital brain functioning. In ostriches, the neck provides an appropriate body part for comparison to the head as it is featherless, reflecting skin temperature, moves in synchrony with the head, and individuals typically orientate themselves such that the side of these two organs can be repeatably captured by thermal imaging. This provides control over distance, angle in the image, and similar skin exposure to sun/shade at the time when being photographed (see Supplementary Information for analyses to support that this approach minimises the effect of distance and angle of imaging on measurements).

## Results

### Thermoregulation across naturally fluctuating temperatures

The surface temperatures (T) of the head and neck were both positively correlated with air temperature (Fig. 2; Tables S1-2). The change in surface temperatures with increasing and decreasing air temperatures was, however, significantly higher for the neck compared to the head (Increasing air T_Neck_ vs Increasing air T_Head_ (CI) = 1.6 (1.2, 1.9), pMCMC = 0.001; Decreasing air T_Neck_ vs Decreasing air T_Head_ (CI) = −2.1 (−3.1, −1.2), pMCMC = 0.001, Fig. 2C; Table S3). Consequently, there were large differences between neck and head temperatures at extreme temperatures (low < 20°C and high > 30°C), but not at benign air temperatures (20 to 30°C), where the need for thermoregulation is minimal (Fig. 2D, Table S4). Furthermore, even though the head and neck differed in their rates of temperature change, they both increased linearly with air temperature (Figs. 2A-B). There was no accelerating change in the surface temperature of the neck (Fig. S1), as would be expected if it acted as a so-called ‘thermal radiator’, directing heat away from the head at critical air temperature inflection points (Tattersall *et al*., 2009; Janse van Vuuren *et al*., 2020). Therefore, head surface temperatures appear to be more tightly regulated than neck surface temperatures at temperature extremes, and there was no evidence that the neck is actively involved in head thermoregulation.

**Fig. 2.**
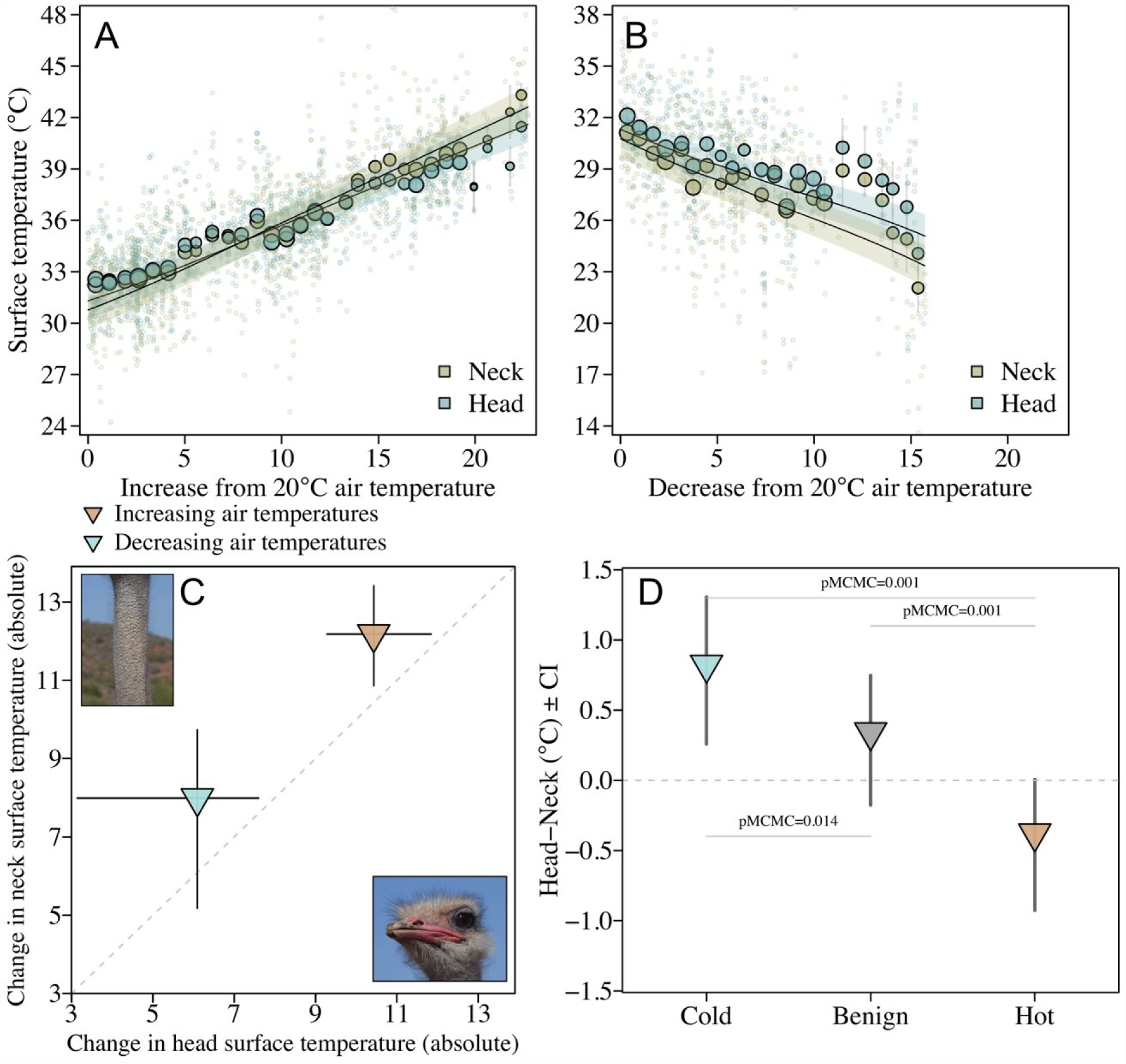
Neck temperatures show how ostriches thermoregulate their head temperatures. **A)** When air temperatures increase above 20°C, the surface temperatures of the head increase slower than the neck (n_images_ = 1895, Tables S1-S2). **B)** Similarly, when air temperatures decrease below 20°C the reduction in surface temperatures are greatest in the neck (n_images_ = 849). Three extreme datapoints in A were removed for graphical purposes (see Fig. S4). The larger points are averages with standard errors binned according to temperature. Point size illustrates relative number of individuals: smallest point = 2 and largest point = 71. Fitted lines and 95% credible intervals (shaded area) were extracted from the statistical models **(C)** The rate of surface temperature change was steeper for the neck compared to the head (Table S3). This difference in rates of temperature increase between the two body parts was consistent across both decreasing and increasing temperatures (n_individuals_ = 423). **(D)** The difference in rates of temperature increase between the head and neck led to higher discrepancy between neck to head surface temperatures during hot conditions (air temperatures >= 30°C, n_images_ = 826) and cold (air temperatures <= 20°C, n_images_ = 849) compared to benign (air temperatures > 20°C & < 30°C, n_images_ = 1069) conditions (Table S4).

### Reproductive consequences and evolutionary potential of head thermoregulation

Next, we examined if the ability to regulate head temperatures under a given thermal load, measured as the difference between head and neck temperatures, has the potential to evolve. We did this by first investigating the relationship of the head-neck temperature with reproductive success and secondly testing if variation across individuals was heritable. We found that females with lower head-neck temperatures during hot periods had higher egg-laying rates (Fig. 3A. Hot_Head-Neck_ (CI) = −0.16 (−0.30, 0.00); pMCMC = 0.027; Table S6). Under more benign temperatures, however, there was no significant relationship between head-neck temperature and egg laying rates (Benign_Head-Neck_ (CI) = −0.06 (−0.18, 0.05); pMCMC = 0.277; Table S6). This suggests that there is selection for the ability to cool the head during high temperatures via its effects on fecundity.

**Fig. 3.**
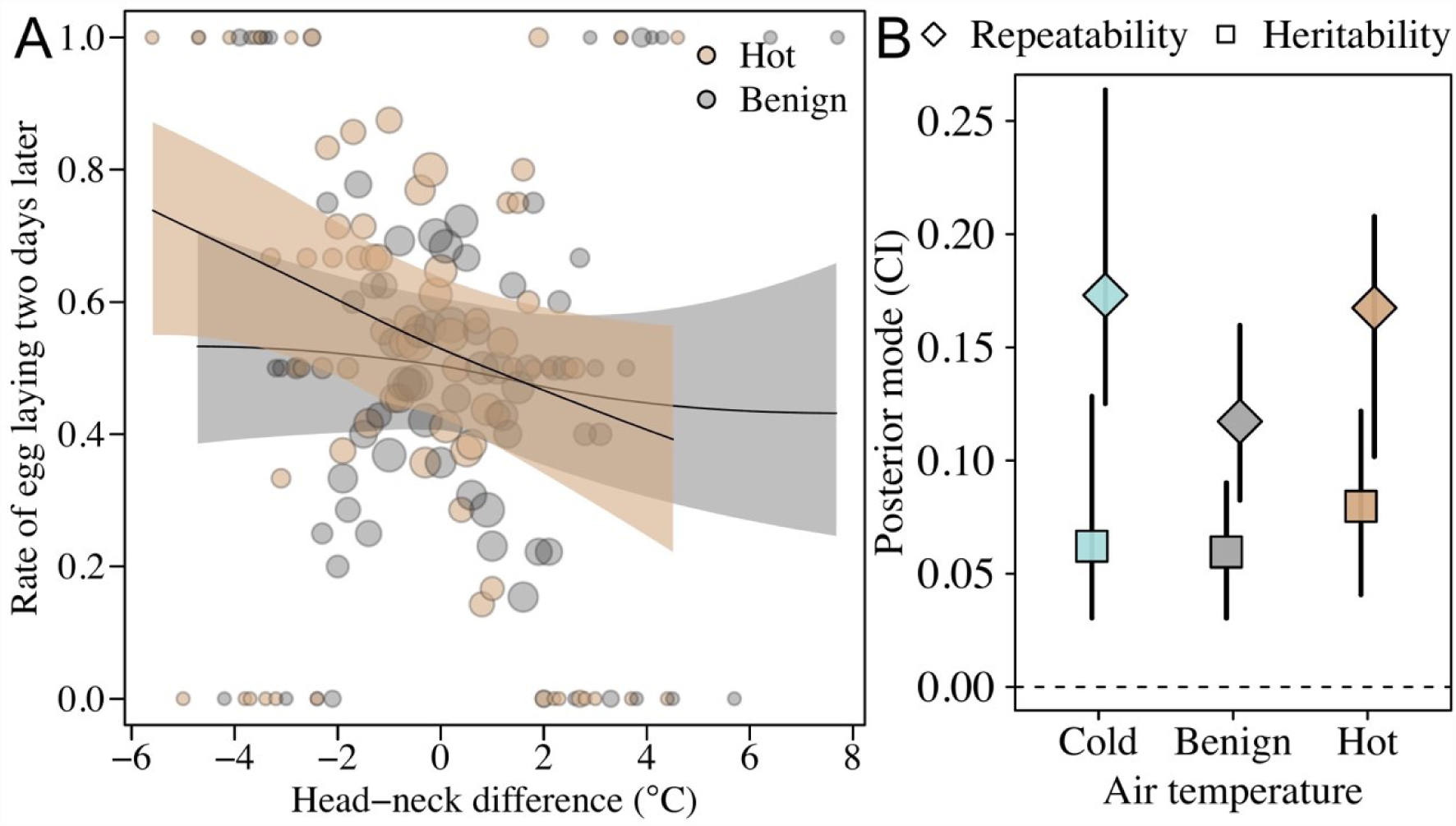
The evolutionary potential for head temperature regulation. The capacity of individuals to regulate their head temperature was estimated as the difference between head and neck temperatures in relation to air temperature. **A)** On hot days (daily maximum > 20°C, n_days_ = 35), when temperatures were highest (air temperatures >= 30°C, n_images_ = 471, n_females_ = 228) an increased capacity to regulate head temperature (i.e. a lower head than neck temperature) was associated with a higher egg-laying rate two days later. There is no such association when examining thermoregulation at times when temperatures were benign of the hot days (air temperatures = 20-30°C, n_images_ = 615, n_females_ = 220) (Table S6). Maximum temperatures below 20°C were recorded only on two days, preventing us from similar analyses on importance of thermoregulation during the cold times of cold days. Fitted lines and 95% credible intervals (shaded area) were extracted from the statistical model **B)** Repeatability and heritability of the head-neck temperature deviation were estimated at different air temperatures measured at the time when thermal images were taken (cold < 20°C, benign = 20-30°C and hot >= 30°C, Table S7). Note that the heritability of these thermal profiles under cold conditions could be estimated because there were cold periods (typically mornings) on days where temperatures exceeded 20°C. This contrasts to selection analyses where temperature per day had to be used because of the two day time lag between egg formation and egg laying.

We further found that head-neck temperature had small but positive and significant heritabilities and evolvabilities (Hansen *et al*., 2011) (h^2^ ranged from 0.06 to 0.08. Fig. 3B, Table S7, see also Table S8-S9 for separate models of head and neck). Heritability estimates were similar and significantly different from zero across cold, benign and hot conditions, although measures of additive genetic variances and repeatabilities were slightly higher under cold and hot conditions. Heritabilities are capped by how repeatable measurements are across individuals, which can often be low for labile physiological traits, such as thermal profiles, when measured non-invasively under semi-natural settings where it is difficult to control for other potentially confounding variables (Figs. 1-2). Our repeatability estimates of the head-neck temperature deviation ranged from 0.12 to 0.17 (Fig. 3B, Table S7). It is therefore plausible that our estimates of heritability, while significant, could underestimate the true heritability of head-neck temperature due to the inherent difficulties in quantifying individual variation in thermoregulatory traits.

### Thermoregulatory ability differs between subspecies from different climatic regions

Finally, we investigated whether head-neck temperature showed some evolutionary signatures in response to past climate conditions by comparing females from different ostrich subspecies that originated from different geographic regions (Fig. 4A-B). At our study site in South Africa, there are three ostrich subspecies from different regions of Africa: South African Blacks (SAB: *Struthio camelus*), Zimbabwean Blues (ZB: *S. c. australis*) and Kenyan Reds (KR: *S. c. massaicus*) that have evolved under different climatic regimes (Fig. 4A-C). The climate in East Africa, where KR naturally occurs, is less seasonal, exhibiting lower fluctuations in temperature and precipitation, than the regions where ZB and SAB subspecies occur (Fig. 4C). Although the number of subspecies is limited, comparing individuals from these three subspecies raised at the same site in South Africa allowed us to examine variation in head-neck temperature in this ‘common garden’ setting.

**Fig. 4.**
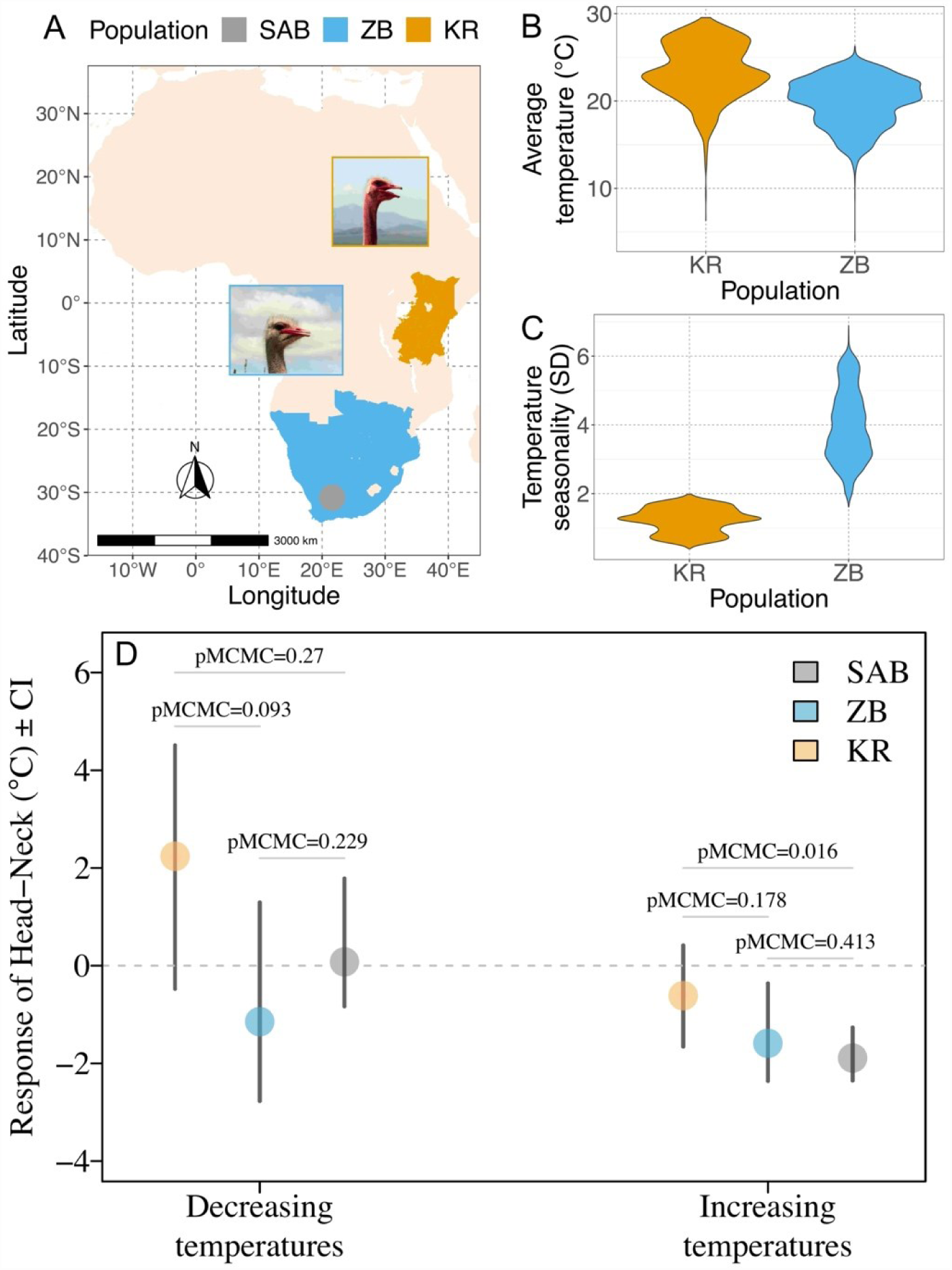
Ostriches originating from more variable climates exhibit larger decreases in head-neck temperatures when hot. **(A-C)** Kenyan Reds (KR) inhabit eastern Africa that is less seasonal and has lower temperature fluctuations compared to Southern Africa where Zimbabwean Blues (ZB) and South African Blacks (SAB) occur. Distribution ranges in A were estimated from regional presence/absence data from Avibase (https://avibase.bsc-eoc.org) and climatic data was obtained from WorldClim (Fick & Hijmans, 2017). Average temperature and seasonality were the main bioclimatic variables of PC1 (43%) and PC2 (29%) in a PCA of 19 bioclimatic variables (Fig. S5 & Table S11). **(D)** For both the SAB (n_individuals_ = 264) and ZB (n_individuals_ = 36) ostriches, the head temperature deviation (head-neck) was reduced when temperatures increased, but this was not the case for KR (n_individuals_ = 23) (Table S12, see also Figs. S6-7 which includes hybrid individuals).

There were pronounced differences in head-neck temperature between the subspecies in response to increasing temperatures that corresponded to the stability of their past climates. The head-neck temperature of KR females, which come from a stable thermal environment, changed little in response to temperature increases (Fig. 4C, Table S12). This indicates similar rates of change in head and neck temperature with air temperature, and therefore only modest thermoregulation in response to increasing temperatures. In contrast, females from ZB and SAB populations, which have their origins in the more variable thermal environments of southern Africa, had head temperatures that did not increase as fast as neck temperatures with rising air temperatures, suggesting a greater capacity to regulate their head temperature (Fig. 4D). In contrast, head-neck temperatures during decreasing temperatures were similar across KRs, SABs and ZABs. These differences in thermoregulatory capacity were not confounded by differences in body size between the three subspecies (Tables S13-14, Figs. S2-3).

## Discussion

Our study has revealed that the capacity to regulate head temperature is important for reproductive success under fluctuating temperatures (Fig. 3). The efficiency of thermoregulation appears to have a genetic basis and enables reproduction under a greater range of temperatures (Fig. 3). Ostriches originating from geographic areas with more pronounced temperature fluctuations were also more efficient at regulating head temperatures, revealed by lower head relative to the neck temperatures. These results suggest that thermoregulation of the head has evolved in response to past climates and may evolve in response to future climatic change (Fig. 4).

Previous research has identified a variety of physiological and behavioural thermoregulatory mechanisms (Kilgore *et al*., 1976; Fuller *et al*., 2003; Huey *et al*., 2003; Angiletta, 2009, 2009; Tattersall *et al*., 2009; Symonds & Tattersall, 2010; Weissenböck *et al*., 2010; Périard *et al*., 2015; Muñoz & Losos, 2018). While such studies have provided compelling evidence for the influence of various traits on thermal physiology, they have not provided direct estimates of genetic variation in such traits or their effect on fitness. Such data is required to fully understand how thermoregulatory traits influence future evolutionary responses to changing climatic conditions. Our results demonstrate the power of combining measurements of reproductive success and pedigree data with thermal profiles from individuals exposed to natural temperature variation in realistic environmental settings. The differences (Fig. 4) between phenotypically different subspecies from different climatic regions in combination with our findings of significant additive genetic variance in head thermoregulation (Fig. 3), suggest that selection for reproducing under fluctuating climates has shaped the evolution of the ostrich’s remarkable capacity for head cooling under hot conditions.

Decades of research on birds in temperate zones has focused on food availability in altricial birds as a major limiting factor for reproduction (Lack, 1954; Williams, 1966). However, for precocial birds inhabiting tropical and subtropical areas, like the ostrich, temperature stress during reproduction might pose a more severe challenge than food limitation (Bourne *et al*., 2020). Thermoregulation can keep body temperatures within non-lethal limits (Tattersall *et al*., 2009; Symonds & Tattersall, 2010; Tattersall & Cadena, 2010; Darnell & Munguia, 2011; McCafferty *et al*., 2011; Szafrańska *et al*., 2020), but heat waves can still compromise population viability by decreasing individual reproductive success (Walsh *et al*., 2019; Bourne *et al*., 2020; Parratt *et al*., 2021). Recent research from different taxa suggests that climate-mediated local extinctions might already be common (Sinervo *et al*., 2010; Wiens, 2016), and there are worrying signs of recent collapses of some desert bird communities (Riddell *et al*., 2019). Whether genetic variation in thermal tolerance is sufficient to drive evolutionary responses to increasingly hot and fluctuating conditions remains an open question.

The low heritability estimates of thermal profiles in this study suggest a low but significant evolutionary potential of ostrich populations to adapt to future temperature change. Previous research on ectotherms, such as insects and reptiles, has also found thermal tolerance to have low additive genetic variances or heritabilities, indicating that obtaining precise quantitative genetic estimates for thermal tolerance may be difficult (Hoffmann *et al*., 2003; Logan *et al*., 2018; Castañeda *et al*., 2019). Variation in body temperatures can also be nonadaptive or maladaptive, a contention supported by recent empirical research (Ghalambor *et al*., 2015; Radersma *et al*., 2020; Stamp & Hadfield, 2020; Svensson *et al*., 2020). For example, plastic changes in either core body temperatures or surface temperatures have been shown to reduce fitness in insects (Svensson *et al*., 2020), reptiles (Campbell-Staton *et al*., 2021), and birds (Nilsson *et al*., 2016). Such maladaptive thermal plasticity may reflect costs of maintaining homeostasis and stable body temperatures under thermally stressful conditions (Svensson *et al*., 2020; Campbell-Staton *et al*., 2021). It is likely that selection for thermal robustness on one side, and costs of maintaining a stable body temperature on the other side, are shaping genetic variance in thermoregulation in both the ostrich and other organisms inhabiting thermally stressful environments like deserts. While challenging, combining studies on the genetics of thermal tolerance with long-term population monitoring of reproduction and survival is key to forecasting the potential effects of climate change on population viability. This is especially the case for large, sub-tropical endotherms like the ostrich.

## Methods

### Study site, subspecies and general settings of enclosures

The study was conducted at the Oudtshoorn Research Farm in the arid Klein Karoo region of South Africa (GPS: 33° 38’ 21.5’S, 22° 15’ 17.4’E). Fenced enclosures (N=170) were used for thermal imaging and to monitor the reproductive success of ostriches in male-female pairs (N_enclosures_ = 148, ~0.25 ha per enclosure) (Cloete *et al*., 2008) and groups (N_enclosures_ = 22, ~0.47 ha per enclosure). These enclosures contained natural vegetation in the form of bushes and small trees, making it possible for the animals to take shelter when needed. All individuals had access to *ad libitum* food and water. The ostriches in this study belong to three different subspecies: 1) the Masai ostrich (*Struthio camelus massaicus*), or Kenyan Red (KR), 2) the Southern African ostrich, (*S. c. australis*), sometimes referred to as the Zimbabwean Blue (ZB) because of its origin in Namibia and Zimbabwe, and 3) the South African Blacks (SAB), that is thought to be of mixed origin, but is genetically similar to ZB (Davies et al 2012; unpublished data). SAB are also referred to as *S. c. var. domesticus*. Individuals that had less than 85% expected relatedness to a particular subspecies, as determined by the pedigree (see below), were considered hybrids. Breeding birds were recruited from surviving chicks from previous years, and parentage data were used to compile a 9-generation pedigree with 139 founding individuals. Although 9 generations have passed since the first individuals were used to establish the study populations, not all individuals are from F9. More than 85% of the photographed ostriches were six years or younger. Ostriches are reproductively mature at age 2-3 years, whereby typically three generations were present in any given year. Unlike the SAB and ZB populations which were founded more than 30 years ago, the KR population was founded just 16 years ago reducing the number of generations across its cohort substantially. Ethical clearance was obtained from the Western Cape Department of Agriculture (DECRA R12/48).

### Thermal imaging data

We took thermal images of ostriches in the enclosures using an infrared thermography camera (H2640, NEC Avio Infrared Technologies; purchased from the company Senso-Test in Sweden). Pictures were taken in the field, usually from distances between 2 m and 25 m. We regularly calibrated the thermal camera against a black body during our field sessions, following the instructions from the camera supplier, which also provided regular recalibration services of our equipment.

Images that were out of focus were discarded prior to analysis, and we used the software InfRec Analyzer to draw separate polygons within the head and neck regions of the ostrich in each image. The average temperature of these polygons was extracted as individual head and neck surface temperatures, respectively. We used the same procedures and default software settings as in our previous work (Svensson & Waller, 2013; Svensson *et al*., 2020), which assumes an emissivity of 1. Biological structures, such as bird skin and feathers, typically have emissivity values ranging between 0.94-1.0 (Gerken *et al*., 2006; Yahav & Giloh, 2012). However, the exact emissivity value used does not change our results as we focused on relative temperature measures that are independent of emissivity values. Hourly air temperatures were measured at a weather station positioned 600 m from the study populations. We fitted a cubic spline to the hourly temperature estimates of each day using the R-package *mgcv* v.1.8 (Wood, 2004), from which we extracted the predicted air temperature at the time-points when thermal images were taken.

### Datasets and analyses

#### 3.1) Thermal imaging dataset

From 2012 to 2017 during October to December, we took 2744 pictures of 423 females between 5 am and 6 pm (average images per female = 6.5). This dataset was designed to maximize the number of individuals with repeated measures within and across years, in order to characterize individual thermal profiles across different thermal environments. Consequently, there was little repeated sampling within days (8% of the pictures).

#### 3.2) General modelling details

We constructed and ran generalized linear mixed models (GLMMs) in R v.3.6.0 (R Core Team, 2020) using the Bayesian framework implemented in the R-package MCMCglmm v.2.29 (Hadfield, 2010). For random terms, we used the weakly informative inverse-Gamma distribution (scale = 0.001, shape = 0.001, i.e. *V = diag(n), nu =n-1+0.002, with n being the dimension of the matrix*) as priors. Unless otherwise stated, each model was run for 5,100,000 iterations of which the initial 100,000 were discarded and only every 4,000th iteration was used for estimating posterior probabilities. The number of iterations was based on the inspection of autocorrelation among posterior samples in preliminary runs. Convergence of the estimates was checked by running the model three times and inspecting the overlap of estimates in trace plots and the level of autocorrelation among posterior samples. We report the results from the first of these three runs. Posterior mode and 95% credible intervals are reported for random effects.

In all analyses, we accounted for environmental effects that varied across years, such as diet, by including year as a random effect. Photographs were taken across 48 days, and we therefore included date as a random effect. We also added enclosure as a random effect as they were repeatedly used across years and varied in vegetation cover, potentially impacting on the local climatic conditions. Individual ID was also included as a random effect, as all analyses contained multiple records per individual.

#### 3.3) Are head surface temperatures more tightly regulated than neck surface temperatures?

To estimate the change in surface temperatures with changing air temperatures we first constructed models with either head or neck temperatures as the Gaussian response variable. Hereafter we ran a model including both head and neck temperatures to allow comparison of the two body parts. We examined increases and decreases in air temperature from a thermal optimum as this allows us to estimate separate parameters for the responses to decreases in temperature (cold tolerance) and increases in temperature (heat tolerance). A previous investigation showed the reproductive success of ostriches is highest at a daily maximum temperature of ~20°C (Schou *et al*., 2021). We therefore defined 20℃ as the optimum air temperature for ostriches, and calculated absolute *temperature change* away from this optimum, with a factor *direction of change* used to denote if it was a decrease or an increase in temperature from the optimum. To make the intercept in statistical models represent the most benign air temperature, we set 20°C to 0 and calculated deviations above (increases) and below (decreases) this value. The variance of slopes (e.g. *temperature change,* see below) depends on the scale of the parameter. We therefore standardized our data by dividing it by the maximum of the air temperature range, resulting in 1 being the maximum temperature change.

Models included the fixed effects of temperature change (ranging from 0 to 1) and direction (decreases or increases). The interaction between temperature change and direction was modelled with a common intercept for decreases and increases, as the way temperature change was calculated dictated that the intercepts were identical. We included the fixed effects of subspecies (SAB, ZB, KR or hybrids) and both the linear and quadratic terms of time of day (scaled and centered to a mean of zero and unit variance). We included interactions between subspecies, temperature change and direction. In addition to the random effects common for all models (year, enclosure, date and individual ID), we also interacted individual ID with temperature change and direction, to allow independent rates of change in surface temperature of each female. This was modelled as a 3×3 unstructured variance-covariance matrix.

The model with both head and neck surface temperatures as response variable contained additional fixed and random effects. To identify the source of the surface temperature we included the fixed effect body part (factorial: neck or head) which was interacted with all the previously described fixed effects. We also added image as an additional random effect because head and neck surface temperatures of an individual at a given time were derived from the same image.

#### 3.4) Does the neck actively regulate the temperature of the head?

To investigate if the neck acts as a thermal radiator actively regulating head temperature, we inspected the relationship between neck surface temperature and air temperature. When air temperature increases towards the body temperature of the ostrich, the difference between air and surface temperature will decrease. If the neck is a thermal radiator, we would expect that as the air temperature reaches certain inflection points, blood is shunted to the surface of the neck to emit more heat, increasing the neck temperature and thereby increasing the difference between neck surface temperatures and air temperature (Tattersall *et al*., 2009; Janse van Vuuren *et al*., 2020). If instead the neck does not function as an actively regulated thermal radiator, we expect a linear relationship between air temperature and neck temperature.

#### 3.5) Effect of head-neck temperature on reproductive success

To test if the capacity to regulate head temperature influences reproductive success we analysed its relationship with rates of egg-laying. Microclimatic factors (i.e. the operative environmental temperature) and the behaviour up to the time of the thermal image being taken will influence the absolute head surface temperatures. To better be able to compare head temperature regulation across individuals, that always will differ in their recent behaviour and microclimatic exposure, we used the neck, which moves in synchrony with the head, as a proxy for the heat stress the animal is experiencing and what it has been exposed to (abiotic and biotic activity). This was done by estimating the head-neck temperature (head surface temperature – neck surface temperature) as a standardised measure of head thermoregulation. Individuals that show lower head-neck temperatures when hot compared to others therefore regulate their head temperature to a larger extent.

After estimating the head-neck temperature of individual females, we connected it to their egg-laying records. From previous investigations we know that when the daily maximum temperature is above or below 20°C, ostrich egg-laying rates start to decrease two to four days later, possibly because this is the time it takes for the egg to travel down the oviduct (Schou *et al*., 2021). The maximum temperature was below 20°C on just two of the days when thermal images were taken, preventing us from examining the relationship between reproduction and temperature regulation under cold conditions. In contrast, maximum temperatures were above 20°C in 41 days when thermal images were taken allowing responses to heat and reproduction to be analysed.

We examined the probability of eggs being laid two to four days after females were thermal imaged on days exceeding 20°C. If females with lower head-neck temperatures have a higher probability of laying an egg during this time window, this would indicate that the thermoregulatory capacity of the head is associated with reproductive success. Any such associations may not be causal, as it is possible that reproduction itself causes an increase in surface temperatures, for example by impacting metabolic rate. Assuming that any impact of reproduction on surface temperatures is similar during hot and benign times of the day, we grouped the data into photographs taken during benign (air temperatures > 20°C & < 30°C, n_images_ = 615) and hot (air temperatures >= 30°C, n_images_ = 471) times of the day. Only if the relationship between head-neck temperature and egg-laying is most pronounced when hot, would we then expect it to be a consequence of improved thermoregulation under heat stress.

The data was analysed using a model with the probability of laying (binary, model type: “threshold”) as the response variable. The fixed effects included were head-neck temperature (scaled and centered to a mean of zero and unit variance), air temperature category (hot or benign), and subspecies (KR, SAB, ZB or Hybrid), as well as the interaction between head-neck temperature and temperature category. As females of two years of age lay fewer eggs than older females (Schou *et al*., 2021), a factor of age (2 versus >2) was also included as a fixed effect. We included year, enclosure, date and individual ID as random effects. The first 45 days of the breeding season were removed as this is the average time it takes for pairs to acclimate to each other and their enclosure (Schou *et al*., 2021). We also removed females that laid fewer than ten eggs per year (5% of all females) to avoid including females from incompatible pairs, and individuals that were not in breeding condition. The models were run for 31,500,000 iterations of which the initial 1,500,000 were discarded and only every 10,000th iteration was used for estimating posterior probabilities.

#### 3.6) Is there genetic variation in the efficiency of head thermoregulation?

To investigate the evolutionary potential of head thermoregulation we constructed models to estimate the repeatability and heritability when cold (air temperatures <= 20°C, n_images_ = 849), benign (air temperatures > 20°C & < 30°C, n_images_ = 1069) and hot (air temperatures >= 30°C, n_images_ = 826). The grouping of air temperature into these three categories was based on the thermal neutral zone of the emu (Maloney, 2008), and to ensure roughly equal replication within the cold and hot categories. Models had head-neck temperature (Gaussian) as the response variable, and we included the fixed effect temperature category (factorial: cold, benign or hot). These categories do not capture the effects of small deviations in air temperature, and we therefore added a continuous measure of the deviation from the mean air temperature in each category. This fixed effect was constructed by centering and scaling the air temperature records within each air temperature category. We also included the fixed effects of subspecies (SAB, ZB, KR or hybrids) and both the linear and quadratic terms of time of day (scaled and centered to a mean of zero and unit variance). We included interactions between subspecies and temperature category. In addition to the random effects common for all models (year, enclosure, date and individual ID), we also interacted individual ID with air temperature category, to allow independent rates of change in surface temperature of each female. This was modelled as a 3×3 unstructured variance-covariance matrix. We also estimated the residual variance separately for each air temperature category.

Repeatability of head-neck temperatures under different temperature categories were estimated using the estimates of permanent individual variances (*pe*) from the variance-covariance matrix of individual ID by air temperature category:

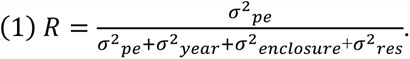

The individual variances and covariances in head-neck temperature may originate from both environmental and genetic factors. To control for maternal effects post laying, eggs are removed from the nest twice a day and artificially incubated. Hereafter chicks were reared in small pens in mixed groups with access to shade, and with *ad libitum* food and water - but without adult ostriches. These experimental procedures should experimentally remove the majority of any maternal effects. We also tried to estimate maternal effects by fitting dam identity in our statistical models. However, such dam effects were close to zero and were therefore excluded from our final models, as they did not significantly improve the explanatory power of our models. This suggests that such maternal effects – even if present – do not persist in to adulthood and are therefore not affecting adult thermal profiles.

To partition the among-individual variance that is due to additive genetic effects we added a second 3×3 unstructured variance-covariance matrix of individual ID linked to the pedigree (*a*). With these variance components, we estimated the narrow sense heritability (h^2^) of head-neck temperature in each air temperature category as the proportion of phenotypic variance attributable to additive genetic variance:

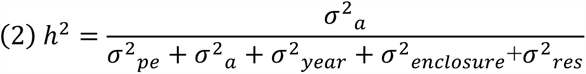

We also estimated evolvability (I_A_) (Houle, 1992):

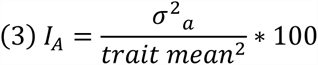

One characteristic of evolvability is that it increases very fast as trait mean approaches zero. When we initially used the posterior of the trait mean to estimate evolvability, this caused near-infinity estimates of evolvability for some of the samples in the posterior, causing biased estimates of the posterior mode and mean of evolvability. To avoid this, we used the posterior mode of trait mean in the denominator, such that only the uncertainty of additive genetic variance is included in the reported estimate of evolvability. We also ran identical models with head or neck surface temperature as the response variable. The outcome of these analyses is available in the supplementary materials (Tables S8-S9). Finally, we also used a similar model to investigate if average feather temperatures on the side of the abdomen better captures temperature exposure of the animal than recorded air temperatures. This model was run without the animal term, and had temperature categories based on individual feather temperature instead of recorded air temperatures. However, temperature measurements of feathers were highly variable (Figure S8), likely because the feathers could have been facing the sun or away from the sun in the time up until the picture. Indeed, estimated repeatabilities of the head-neck difference was not higher when using feather temperatures instead of air temperatures (Table S5).

#### 3.7) Current distribution of the ostrich subspecies

The different ostrich subspecies come from different regions of Africa that may differ in climate and therefore impact the need for thermoregulation. We obtained estimates of the current distributions of *S. c. massaicus* (KR) and *S. c. australis* (ZB) were obtained by downloading region-based presence/absence data from Avibase (https://avibase.bsc-eoc.org, September, 2020) and plotting these using the R-package “rnaturalearth” v. 0.1.0. To characterise the climate of the distributions of each subspecies, we downloaded 19 bioclimatic variables (10min) from WorldClim v. 2 (Fick & Hijmans, 2017). We performed a Principal Component Analyses (PCA) and inspected the loadings of the first four principal components after varimax transformation (Table S11). Based on this inspection we described each principal component by one or two bioclimatic variables to characterize climatic differences in among the distributions of the ostrich subspecies.

#### 3.8) Do ostriches originating from different climatic regions differ in thermoregulation?

To test if the subspecies differ in their responses to increasing and decreasing temperatures, we modelled head-neck temperature in a random regression model. This approach also allows us to test if the head-neck temperature at the optimum temperature (the intercept) influences the rate of change in head-neck temperatures as air temperatures increase or decrease (the slopes), and if such a relationship is genetically based. Models included the fixed effects of temperature change (ranging from 0 to 1) and direction (decreases or increases). The interaction between temperature change and direction was modelled with a common intercept for decreases and increases, as the way temperature change was calculated dictated that the intercepts were identical. We included the fixed effects of subspecies (SAB, ZB, KR or hybrids) and both the linear and quadratic terms of time of day (scaled and centered to a mean of zero and unit variance). We included interactions between subspecies, temperature change and direction. In addition to the random effects common for all models (year, enclosure, date and individual ID), we also interacted individual ID with temperature change and direction, to allow independent rates of change in surface temperature of each female. This was modelled as a 3×3 unstructured variance-covariance matrix. To partition genetic and non-genetic effects contributing to the relationship between temperature at the optimum and temperature change, we added a second 3×3 unstructured variance-covariance matrix of individual ID linked to the pedigree and interacted with temperature change and direction. The genetic variance and co-variance was then used to estimate the genetic correlation between the slopes and intercepts (correlation = covariance_trait1,trait2_ / sqrt(var_trait1_*var_trait2_)) (Table S10).

To test if differences in the surface temperatures of the head and neck under increasing or decreasing air temperatures across subspecies were caused by differences in body mass, we ran a set of identical models also including individual body mass (scaled and centered to a mean of zero and unit variance) as a fixed effect. We had records of body mass for all individuals and when multiple records were available for one individual, we used the record closest to the time of the thermal image.

#### 3.9) The effects of distance and angle on surface temperatures

Animal surface temperatures measured using thermal imaging can be influenced by confounding factors such as distance to the animal and animal orientation to the sun and wind speed. We quantified the impact of distance to the animal on estimated surface temperatures, using the number of pixels in the head and neck region as a proxy for distance. When we inspected the relationship between number of pixels and absolute head or neck surface temperatures, we found indications of more error in images where birds are further away. However, in pictures taken far away, both head and neck are small, resulting in any biases being accounted for by the head-neck metric (Fig S9). The angle of the animal is also important, and ostriches typically orientate themselves side on to people so that the side of the head and neck consistently face the investigator. This ensures that the majority of pictures are taken from the same angle. To quantify this, we recorded the angle of the head in a subset of pictures and found 87% included the side of the head, and found no effect of the angle of the head on the head-neck metric (Fig S10). In conclusion, our measures of head-neck temperature appear robust to confounding variables such as distance and angle. This is in line with several independent studies that have shown to surface temperatures to correlate with internal body temperatures (McCafferty *et al*., 2015; Andreasson *et al*., 2016; Szafrańska *et al*., 2020; Gauchet *et al*., 2022) and that the effect of confounding variables on measurements of internal body temperatures are relatively modest (Gauchet *et al*., 2022). Finally, additional details about our statistical analyses of thermal imaging data and results are provided in the Supplementary Materials (Tables S1-S16 and Figs. S1-S10).

## Acknowledgements

We thank the staff and workers at Oudtshoorn Research Farm for assistance with data collection and maintenance of the birds and the Western Cape Government for use of their resources. We also thank members of “Svensson Lab” (students and assistants) who helped with analyzing the thermal images, and Andreas Nord and Fredrik Andreasson for valuable feedback on an earlier version of this manuscript. The computations were performed on resources provided by SNIC through Uppsala Multidisciplinary Center for Advanced Computational Science (UPPMAX) under Project SNIC 2018/8-359.

## Author Contribution

Conceptualization: E.I.S, C.K.S., M.F.S.; Data curation: E.I.S., C.K.C., M.F.S., A.E., Z.B., S.C.; Formal analysis: M.F.S.; Funding acquisition: C.K.C, E.I.S., M.F.S, S.C.; Investigation: E.I.S., C.K.C., J.M., J.W., M.F.S., S.C., Z.B; Methodology: E.I.S., C.K.C., M.F.S.; Project administration: E.I.S., C.K.C., M.F.S.; Writing-original draft E.I.S, C.K.C., M.F.S.; Writing – reviewing and editing: A.E., CKC, E.I.S., J.M., J.W., M.F.S, S.C., Z.B.

## Funding

Carlsberg Foundation (MFS)

Swedish Research Council grant 2017-03880 (CKC)

Knut and Alice Wallenberg Foundation grant 2018.0138 (CKC)

Carl Tryggers grant 12: 92 & 19: 71 (CKC)

Carl Tryggers grant 19: 71 (CKC)

Swedish Research Council grant 2016-03356 (EIS)

Western Cape Agricultural Research Trust grant 0070/000VOLSTRUISE (SC)

Technology and Human Resources for Industry program of the South African National

Research Foundation grant TP14081390585) (SC)

## Data Accessibility

All data extracted from thermal images are available at https://osf.io/fu2wx/ and will be uploaded in an open digital repository such as Dryad Digital Repository (https://datadryad.org/stash) upon acceptance on this manuscript. The remaining data used support the findings of this study are available from the Western Cape Department of Agriculture in South Africa (WCDA). Restrictions apply to the use of some of these data, which are thus not publicly available. These data are however available from the WCDA upon reasonable request. Code for analyses is available on Github: www.github.com/abumadsen/thermal-images-ostrich/tree/main

## Competing interests

Authors declare that they have no competing interests.

## Supporting Information

Tables S1-S16 Figures S1-S10

## Notes

https://osf.io/fu2wx/

### Competing Interest Statement

The authors have declared no competing interest.

### Summary of Updates

This manuscript has been updated to include additional analyses, following the suggestions and input from an Associate Editor and three referees.

