## Supplementary Material for "Heritable variation in thermoregulation is associated with reproductive success in the world’s largest bird"

Figure S5: Four first principal components from PCA on bioclimatic variables in the ostrich range 22

Figure S6: Change in relative head temperature with increasing air temperature across subspecies 23

Figure S7: Change in relative head temperature with decreasing air temperature across subspecies 24

### Supplementary Tables

#### Glossary of model terms

##### General terms

- SubSpe = Subspecies (factorial: SAB, ZB, KR or Hybrid)
- Daytime_z = Day time (continuous, hours), scaled and centered
- Time = Time during day (factorial: Morning or Afternoon)
- Daytime_ztime = Day time (continuous, hours), scaled and centered within the Time term
- BodyPart = Body part (factorial: Head or Neck)
- Mass_z = Body mass (continuous, kg), scaled and centered
- HeadNeckDiff_z = Difference in surface temperature between head and neck on each thermal image (continuous, °C), scaled and centered
- Age = Age (factorial, >=2 years old[yo] or <2 years old)

##### Terms specific to random regression models of air temperature

- TempCon = Absolute air temperature change from optimum (continuous)
- TempDir = Direction of air temperature change (factorial: decreasing from optimum [Dec] or increasing from optimum [Inc])

##### Terms specific to character state models of air temperature

- TempCat = Air temperature category (factorial: Benign, Cold or Hot)
- Temp_zcat = Air temperature, scaled and centered within the TempCat term
- MaxTempCat = Maximum daily temperature category (factorial: Benign or Hot)

#### Thermal plasticity (Supplementary tables 1-5)

**Table S1: Results from model of the change in head surface temperature with increasing/decreasing temperatures (random regression).**

| Type | Term | Estimate (CI) | pMCMC | Level |
|---|---|---|---|---|
| **Fixed effects** | Intercept | 30.97 (29.96,31.86) | **0.001** | - |
| | SubSpe(ZB) | -0.43 (-0.98,0.15) | 0.131 | - |
| | SubSpe(KR) | 0.09 (-0.63,0.76) | 0.792 | - |
| | SubSpe(Hybrid) | 0 (-0.46,0.36) | 0.986 | - |
| | poly(Daytime_z, 2)1 | 12.68 (-1.28,25.33) | 0.062 | - |
| | poly(Daytime_z, 2)2 | -26.74 (-34.29,-18.24) | **0.001** | - |
| | TempCon:TempDir(Dec) | -6.12 (-8.33,-3.92) | **0.001** | - |
| | TempCon:TempDir(Inc) | 10.34 (9.13,11.57) | **0.001** | - |
| | TempCon:TempDir(Dec):SubSpe(ZB) | -1.32 (-4.11,1.27) | 0.285 | - |
| | TempCon:TempDir(Inc):SubSpe(ZB) | -0.12 (-1.52,1.08) | 0.878 | - |
| | TempCon:TempDir(Dec):SubSpe(KR) | -0.59 (-3.8,2.77) | 0.718 | - |
| | TempCon:TempDir(Inc):SubSpe(KR) | -1.39 (-2.59,0.15) | 0.053 | - |
| | TempCon:TempDir(Dec):SubSpe(Hybrid) | -0.31 (-2.44,1.61) | 0.747 | - |
| | TempCon:TempDir(Inc):SubSpe(Hybrid) | -0.61 (-1.52,0.28) | 0.198 | - |
| **Random effects** | (Intercept) | 0.356 (0.176,0.659) | - | ID |
| | TempCon:TempDir(Dec) | 7.457 (2.072,15.392) | - | ID |
| | TempCon:TempDir(Inc) | 0.526 (0.221,1.611) | - | ID |
| | enclosure | 0.074 (0.015,0.196) | - | enclosure |
| | date | 0.587 (0.346,1.128) | - | date |
| | year | 0.35 (0.007,3.045) | - | year |
| | residuals | 4.159 (3.88,4.409) | - | residuals |
| **Correlations** | TempCon:TempDir(Dec):(Intercept).ID | -0.12 (-0.5,0.57) | 0.939 | - |
| | TempCon:TempDir(Inc):(Intercept).ID | -0.67 (-0.9,-0.3) | **0.01** | - |
| | TempCon:TempDir(Inc):TempCon:TempDir(Dec).ID | -0.02 (-0.8,0.52) | 0.682 | - |

Estimate: Posterior mean is used for fixed effects and posterior mode is used for random effects

**Table S2: Results from model of the change in neck surface temperature with increasing/decreasing temperatures (random regression).**

| Type | Term | Estimate (CI) | pMCMC | Level |
|---|---|---|---|---|
| Fixed effects | Intercept | 30.33 (29.26,31.23) | **0.001** | - |
| | SubSpe(ZB) | -0.33 (-0.95,0.35) | 0.304 | - |
| | SubSpe(KR) | 0.7 (-0.11,1.46) | 0.082 | - |
| | SubSpe(Hybrid) | 0.13 (-0.3,0.62) | 0.606 | - |
| | poly(Daytime_z, 2)1 | 19.93 (3.99,34.91) | **0.008** | - |
| | poly(Daytime_z, 2)2 | -31.2 (-40.33,-22.11) | **0.001** | - |
| | TempCon:TempDir(Dec) | -6.55 (-9.1,-3.97) | **0.001** | - |
| | TempCon:TempDir(Inc) | 12.03 (10.62,13.46) | **0.001** | - |
| | TempCon:TempDir(Dec):SubSpe(ZB) | -0.11 (-3.03,2.94) | 0.939 | - |
| | TempCon:TempDir(Inc):SubSpe(ZB) | -0.2 (-1.64,1.27) | 0.819 | - |
| | TempCon:TempDir(Dec):SubSpe(KR) | -2.11 (-5.99,1.59) | 0.286 | - |
| | TempCon:TempDir(Inc):SubSpe(KR) | -2.48 (-3.93,-1.02) | **0.001** | - |
| | TempCon:TempDir(Dec):SubSpe(Hybrid) | -0.96 (-3.29,1.34) | 0.384 | - |
| | TempCon:TempDir(Inc):SubSpe(Hybrid) | -0.7 (-1.65,0.35) | 0.202 | - |
| Random effects | (Intercept) | 0.383 (0.195,0.786) | - | ID |
| | TempCon:TempDir(Dec) | 15.146 (5.747,24.7) | - | ID |
| | TempCon:TempDir(Inc) | 0.482 (0.153,1.506) | - | ID |
| | enclosure | 0.166 (0.05,0.294) | - | enclosure |
| | date | 0.77 (0.416,1.421) | - | date |
| | year | 0.195 (0.001,2.823) | - | year |
| | residuals | 5.276 (4.892,5.554) | - | residuals |
| Correlations | TempCon:TempDir(Dec):(Intercept).ID | 0.03 (-0.48,0.56) | 0.878 | - |
| | TempCon:TempDir(Inc):(Intercept).ID | -0.7 (-0.88,-0.23) | **0.013** | - |
| | TempCon:TempDir(Inc):TempCon:TempDir(Dec).ID | -0.14 (-0.77,0.57) | 0.73 | - |

Estimate: Posterior mean is used for fixed effects and posterior mode is used for random effects

**Table S3: Results from model of the change in surface temperature (neck and head) with increasing/decreasing temperatures (random regression).**

In this model surface temperature was used as a response variable, and the fixed effect *bodypart* was included to differentiate between surface temperatures originating from the neck or head.

| Type | Term | Estimate (CI) | pMCMC | Level |
|---|---|---|---|---|
| Fixed effects | BodyPartNeckAve | 30.37 (29.32,31.23) | **0.001** | - |
| | BodyPart(Head) | 30.88 (29.83,31.74) | **0.001** | - |
| | SubSpe(ZB) | -0.42 (-1.01,0.17) | 0.16 | - |
| | SubSpe(KR) | 0.61 (-0.04,1.41) | 0.094 | - |
| | SubSpe(Hybrid) | 0.14 (-0.29,0.62) | 0.563 | - |
| | poly(Daytime_z, 2)1 | 18.28 (-1.25,38.53) | 0.074 | - |
| | poly(Daytime_z, 2)2 | -38.89 (-49.75,-26.77) | **0.001** | - |
| | TempCon:TempDir(Dec) | -7.58 (-9.73,-5.18) | **0.001** | - |
| | TempCon:TempDir(Inc) | 12.07 (10.86,13.41) | **0.001** | - |
| | BodyPart(Head):SubSpe(ZB) | 0.04 (-0.29,0.38) | 0.813 | - |
| | BodyPart(Head):SubSpe(KR) | -0.53 (-0.97,-0.13) | **0.016** | - |
| | BodyPart(Head):SubSpe(Hybrid) | -0.15 (-0.45,0.11) | 0.274 | - |
| | BodyPart(Head):poly(Daytime_z, 2)1 | 4.41 (-6.1,13.2) | 0.358 | - |
| | BodyPart(Head):poly(Daytime_z, 2)2 | -1.84 (-7.83,3.86) | 0.55 | - |
| | TempCon:TempDir(Dec):SubSpe(ZB) | 0.18 (-2.67,2.91) | 0.89 | - |
| | TempCon:TempDir(Inc):SubSpe(ZB) | -0.34 (-1.59,1.16) | 0.634 | - |
| | TempCon:TempDir(Dec):SubSpe(KR) | -1.53 (-5.04,1.75) | 0.381 | - |
| | TempCon:TempDir(Inc):SubSpe(KR) | -2.48 (-3.89,-1.01) | **0.001** | - |
| | TempCon:TempDir(Dec):SubSpe(Hybrid) | -0.96 (-3.15,0.95) | 0.36 | - |
| | TempCon:TempDir(Inc):SubSpe(Hybrid) | -0.91 (-1.92,-0.01) | 0.056 | - |
| | BodyPart(Head):TempCon:TempDir(Dec) | 2.06 (1.17,3.07) | **0.001** | - |
| | BodyPart(Head):TempCon:TempDir(Inc) | -1.55 (-1.85,-1.23) | **0.001** | - |
| | BodyPart(Head):TempCon:TempDir(Dec):SubSpe(ZB) | -1.58 (-3.18,-0.07) | **0.048** | - |
| | BodyPart(Head):TempCon:TempDir(Inc):SubSpe(ZB) | 0.22 (-0.61,1.18) | 0.627 | - |
| | BodyPart(Head):TempCon:TempDir(Dec):SubSpe(KR) | 1.02 (-0.79,3.05) | 0.318 | - |
| | BodyPart(Head):TempCon:TempDir(Inc):SubSpe(KR) | 1.15 (0.23,2.1) | **0.026** | - |
| | BodyPart(Head):TempCon:TempDir(Dec):SubSpe(Hybrid) | 0.96 (-0.23,2.04) | 0.099 | - |
| | BodyPart(Head):TempCon:TempDir(Inc):SubSpe(Hybrid) | 0.39 (-0.2,1.03) | 0.227 | - |
| Random effects | (Intercept) | 0.357 (0.189,0.717) | - | ID |
| | TempCon:TempDir(Dec) | 7.122 (2.509,15.769) | - | ID |
| | TempCon:TempDir(Inc) | 0.48 (0.174,1.38) | - | ID |
| | Image | 3.499 (3.256,3.795) | - | Image |
| | enclosure | 0.107 (0.034,0.222) | - | enclosure |
| | date | 0.754 (0.416,1.293) | - | date |
| | year | 0.379 (0.001,2.817) | - | year |
| | residuals | 1.321 (1.273,1.409) | - | residuals |

Estimate: Posterior mean is used for fixed effects and posterior mode is used for random effects

**Table S4: Results from model of the difference between head and neck surface temperature across cold, benign or hot air temperatures (character-state)**

In this model the difference between head and neck temperature (head-neck) at each thermal image was used as the response variable.

| Type | Term | Estimate (CI) | pMCMC | Level |
|---|---|---|---|---|
| Fixed effects | Intercept | 0.28 (-0.18,0.75) | 0.202 | - |
| | Temp_zcat | -0.11 (-0.26,0.02) | 0.107 | - |
| | TempCat(Cold) | 0.52 (0.12,0.89) | **0.014** | - |
| | TempCat(Hot) | -0.74 (-0.98,-0.44) | **0.001** | - |
| | poly(Daytime_z, 2)1 | -4.75 (-13.62,4.63) | 0.288 | - |
| | poly(Daytime_z, 2)2 | 2.54 (-2.83,7.51) | 0.355 | - |
| | SubSpe(ZB) | -0.17 (-0.56,0.16) | 0.342 | - |
| | SubSpe(KR) | -0.47 (-0.93,-0.04) | **0.034** | - |
| | SubSpe(Hybrid) | -0.06 (-0.3,0.2) | 0.632 | - |
| | Temp_zcat:TempCat(Cold) | 0.04 (-0.24,0.29) | 0.786 | - |
| | Temp_zcat:TempCat(Hot) | -0.28 (-0.49,-0.06) | **0.013** | - |
| | TempCat(Cold):SubSpe(ZB) | -0.16 (-0.76,0.48) | 0.59 | - |
| | TempCat(Hot):SubSpe(ZB) | 0.42 (-0.08,1.03) | 0.123 | - |
| | TempCat(Cold):SubSpe(KR) | 0.16 (-0.56,0.94) | 0.677 | - |
| | TempCat(Hot):SubSpe(KR) | 0.6 (-0.01,1.14) | 0.053 | - |
| | TempCat(Cold):SubSpe(Hybrid) | 0 (-0.49,0.43) | 0.97 | - |
| | TempCat(Hot):SubSpe(Hybrid) | 0.04 (-0.31,0.39) | 0.835 | - |
| Random effects | TempCat(Benign) | 0.17 (0.103,0.266) | - | ID |
| | TempCat(Cold) | 0.551 (0.33,0.853) | - | ID |
| | TempCat(Hot) | 0.211 (0.123,0.322) | - | ID |
| | year | 0.109 (0.04,0.89) | - | year |
| | enclosure | 0.001 (0,0.026) | - | enclosure |
| | date | 0.044 (0.004,0.088) | - | date |
| | TempCat(Benign) | 2.08 (1.844,2.215) | - | residuals |
| | TempCat(Cold) | 2.921 (2.567,3.213) | - | residuals |
| | TempCat(Hot) | 1.631 (1.461,1.802) | - | residuals |
| Fixed effect contrasts | TempCat(Cold) vs TempCat(Hot) | 1.27 (0.79,1.68) | **0.001** | - |
| Repeatabilities | Hot | 0.095 (0.053,0.153) | - | - |
| | Benign | 0.062 (0.037,0.109) | - | - |
| | Cold | 0.158 (0.085,0.221) | - | - |

Estimate: Posterior mean is used for fixed effects and posterior mode is used for random effects

**Table S5: Results from model of the difference between head and neck surface temperature across cold, benign or hot feather temperatures (character-state).**

In this model we used feather temperatures of the side of abdomen as a measure of temperature exposure instead of air temperature. The difference between head and neck temperature at each thermal image was used a the response variable.

| Type | Term | Estimate (CI) | pMCMC | Level |
|---|---|---|---|---|
| **Fixed effects** | Intercept | 0.18 (-0.45,0.78) | 0.462 | - |
| | BodyAve_z | -0.14 (-0.23,-0.04) | **0.008** | - |
| | BodyAveCatCold | 0.44 (0.18,0.7) | **0.001** | - |
| | BodyAveCatHot | -0.46 (-0.69,-0.24) | **0.001** | - |
| | poly(Daytime_z, 2)1 | -10.2 (-15,-5.39) | **0.001** | - |
| | poly(Daytime_z, 2)2 | -1.04 (-5.33,3.05) | 0.611 | - |
| | SubSpe(ZB) | -0.08 (-0.42,0.27) | 0.634 | - |
| | SubSpe(KR) | -0.4 (-0.86,0) | 0.069 | - |
| | SubSpe(Hybrid) | -0.12 (-0.36,0.17) | 0.382 | - |
| | BodyAve_z:BodyAveCatCold | -0.08 (-0.25,0.1) | 0.368 | - |
| | BodyAve_z:BodyAveCatHot | -0.18 (-0.33,-0.02) | **0.026** | - |
| | BodyAveCatCold:SubSpe(ZB) | -0.33 (-0.96,0.25) | 0.307 | - |
| | BodyAveCatHot:SubSpe(ZB) | 0.36 (-0.2,1.02) | 0.264 | - |
| | BodyAveCatCold:SubSpe(KR) | -0.05 (-0.87,0.62) | 0.926 | - |
| | BodyAveCatHot:SubSpe(KR) | 0.51 (-0.23,1.16) | 0.162 | - |
| | BodyAveCatCold:SubSpe(Hybrid) | 0.11 (-0.3,0.56) | 0.616 | - |
| | BodyAveCatHot:SubSpe(Hybrid) | -0.01 (-0.43,0.39) | 0.974 | - |
| **Random effects** | BodyAveCatBenign | 0.206 (0.119,0.369) | - | ID |
| | BodyAveCatCold | 0.359 (0.184,0.554) | - | ID |
| | BodyAveCatHot | 0.282 (0.126,0.388) | - | ID |
| | year | 0.183 (0.039,1.386) | - | year |
| | enclosure | 0.001 (0,0.021) | - | enclosure |
| | date | 0.02 (0,0.075) | - | date |
| | BodyAveCatBenign | 1.918 (1.742,2.172) | - | residuals |
| | BodyAveCatCold | 2.782 (2.415,3.008) | - | residuals |
| | BodyAveCatHot | 2.036 (1.86,2.324) | - | residuals |
| **Fixed effect contrasts** | BodyAveCatCold vs BodyAveCatHot | 0.95 (0.59,1.21) | **0.001** | - |
| **Repeatabilities** | Hot | 0.088 (0.033,0.143) | - | - |
| | Benign | 0.087 (0.036,0.145) | - | - |
| | Cold | 0.106 (0.049,0.159) | - | - |

Estimate: Posterior mean is used for fixed effects and posterior mode is used for random effects

#### Impact of surface temperature on reproductive success (Supplementary table 6)

**Table S6: Results from analyses of impact of temperature differences between head and neck on egg-laying rate (character-state).**

| **Type** | **Term** | **Estimate (CI)** | **pMCMC** | **Level** |
|---|---|---|---|---|
| **Fixed effects** | TempCat(Benign) | 0.14 (-0.18,0.45) | 0.367 | - |
| | TempCat(Hot) | 0.25 (-0.07,0.59) | 0.113 | - |
| | SubSpe(KR) | -0.39 (-0.91,0.05) | 0.103 | - |
| | SubSpe(SAB) | -0.11 (-0.37,0.18) | 0.424 | - |
| | SubSpe(ZB) | -0.43 (-0.84,0) | **0.043** | - |
| | Age(>=2yo) | -0.03 (-0.28,0.2) | 0.842 | - |
| | TempCat(Benign):HeadNeckDiff_z | -0.06 (-0.18,0.05) | 0.277 | - |
| | TempCat(Hot):HeadNeckDiff_z | -0.16 (-0.3,0) | **0.027** | - |
| **Random effects** | ID | 0.387 (0.2,0.65) | - | ID |
| | year | 0.005 (0,0.163) | - | year |
| | enclosure | 0.002 (0,0.21) | - | enclosure |
| | date | 0.057 (0.001,0.138) | - | date |
| | residuals | 1 (1,1) | - | residuals |

Estimate: Posterior mean is used for fixed effects and posterior mode is used for random effects. The term TempCat(Hot):HeadNeckDiff_z represent

#### Evolutionary potential of surface temperatures (Supplementary tables 7-10)

**Table S7: Results from animal model of head-neck differences in surface temperature (character-state).**

In this model the difference between head and neck temperature at each thermal image was used a the response variable.

| Type | Term | Estimate (CI) | pMCMC | Level |
|---|---|---|---|---|
| **Fixed effects** | Intercept | 0.31 (-0.19,0.75) | 0.168 | - |
| | Temp_zcat | -0.11 (-0.23,0.02) | 0.102 | - |
| | TempCat(Cold) | 0.52 (0.04,0.96) | **0.03** | - |
| | TempCat(Hot) | -0.77 (-1.11,-0.43) | **0.002** | - |
| | poly(Daytime_z, 2)1 | -4.35 (-13.74,3.79) | 0.342 | - |
| | poly(Daytime_z, 2)2 | 2.12 (-3.16,7.25) | 0.416 | - |
| | SubSpe(ZB) | -0.26 (-0.69,0.17) | 0.242 | - |
| | SubSpe(KR) | -0.52 (-1.06,0.09) | 0.078 | - |
| | SubSpe(Hybrid) | -0.13 (-0.43,0.17) | 0.432 | - |
| | Temp_zcat:TempCat(Cold) | 0.04 (-0.21,0.32) | 0.744 | - |
| | Temp_zcat:TempCat(Hot) | -0.27 (-0.48,-0.07) | **0.008** | - |
| | TempCat(Cold):SubSpe(ZB) | -0.06 (-0.8,0.74) | 0.893 | - |
| | TempCat(Hot):SubSpe(ZB) | 0.54 (-0.06,1.25) | 0.096 | - |
| | TempCat(Cold):SubSpe(KR) | 0.22 (-0.71,1.21) | 0.653 | - |
| | TempCat(Hot):SubSpe(KR) | 0.59 (-0.16,1.37) | 0.126 | - |
| | TempCat(Cold):SubSpe(Hybrid) | 0.1 (-0.4,0.7) | 0.718 | - |
| | TempCat(Hot):SubSpe(Hybrid) | 0.08 (-0.33,0.51) | 0.685 | - |
| **Random effects** | TempCat(Benign) | 0.149 (0.092,0.235) | - | ID |
| | TempCat(Cold) | 0.453 (0.188,0.716) | - | ID |
| | TempCat(Hot) | 0.158 (0.096,0.263) | - | ID |
| | TempCat(Benign) | 0.151 (0.081,0.236) | - | animal |
| | TempCat(Cold) | 0.226 (0.115,0.512) | - | animal |
| | TempCat(Hot) | 0.186 (0.089,0.268) | - | animal |
| | year | 0.122 (0.027,0.71) | - | year |
| | enclosure | 0.002 (0,0.026) | - | enclosure |
| | date | 0.028 (0.002,0.079) | - | date |
| | TempCat(Benign) | 2 (1.83,2.191) | - | residuals |
| | TempCat(Cold) | 2.904 (2.55,3.207) | - | residuals |
| | TempCat(Hot) | 1.606 (1.408,1.752) | - | residuals |
| **Repeatabilities** | Hot | 0.167 (0.102,0.208) | - | - |
| | Benign | 0.117 (0.082,0.16) | - | - |
| | Cold | 0.173 (0.125,0.264) | - | - |
| **Heritabilities** | Hot | 0.08 (0.041,0.122) | - | - |
| | Benign | 0.06 (0.03,0.09) | - | - |
| | Cold | 0.062 (0.03,0.128) | - | - |
| **Evolvabilities** | Hot | 150.675 (72.002,217.199) | - | - |
| | Benign | 247.12 (133.592,386.818) | - | - |
| | Cold | 26.965 (13.791,61.17) | - | - |

Estimate: Posterior mean is used for fixed effects and posterior mode is used for random effects

**Table S8: Results from animal model of head surface temperature (character-state).**

| **Type** | **Term** | **Estimate (CI)** | **pMCMC** | **Level** |
|---|---|---|---|---|
| **Fixed effects** | Intercept | 33.17 (32.09,34.18) | **0.001** | - |
| | Temp_zcat | 1.22 (0.95,1.47) | **0.001** | - |
| | TempCat(Cold) | -3.75 (-4.5,-2.94) | **0.001** | - |
| | TempCat(Hot) | 4.42 (3.8,5.09) | **0.001** | - |
| | poly(Daytime_z, 2)1 | 9.43 (-6.32,25.15) | 0.238 | - |
| | poly(Daytime_z, 2)2 | -25.38 (-33.79,-16.7) | **0.001** | - |
| | SubSpe(ZB) | -0.81 (-1.46,-0.27) | **0.018** | - |
| | SubSpe(KR) | -0.02 (-0.77,0.76) | 0.968 | - |
| | SubSpe(Hybrid) | -0.19 (-0.64,0.25) | 0.413 | - |
| | Temp_zcat:TempCat(Cold) | -0.26 (-0.71,0.2) | 0.259 | - |
| | Temp_zcat:TempCat(Hot) | 0.56 (0.09,0.97) | **0.008** | - |
| | TempCat(Cold):SubSpe(ZB) | 0.14 (-0.72,1.09) | 0.784 | - |
| | TempCat(Hot):SubSpe(ZB) | 0.63 (-0.27,1.56) | 0.186 | - |
| | TempCat(Cold):SubSpe(KR) | -0.21 (-1.27,0.93) | 0.741 | - |
| | TempCat(Hot):SubSpe(KR) | -0.88 (-2.12,0.06) | 0.094 | - |
| | TempCat(Cold):SubSpe(Hybrid) | -0.02 (-0.68,0.64) | 0.944 | - |
| | TempCat(Hot):SubSpe(Hybrid) | -0.24 (-0.82,0.36) | 0.43 | - |
| **Random effects** | TempCat(Benign) | 0.31 (0.147,0.538) | - | ID |
| | TempCat(Cold) | 0.323 (0.163,0.672) | - | ID |
| | TempCat(Hot) | 0.197 (0.108,0.401) | - | ID |
| | TempCat(Benign) | 0.293 (0.135,0.5) | - | animal |
| | TempCat(Cold) | 0.235 (0.148,0.58) | - | animal |
| | TempCat(Hot) | 0.234 (0.116,0.39) | - | animal |
| | year | 0.42 (0.006,3.319) | - | year |
| | enclosure | 0.061 (0.014,0.187) | - | enclosure |
| | date | 0.634 (0.33,1.09) | - | date |
| | TempCat(Benign) | 3.808 (3.513,4.238) | - | residuals |
| | TempCat(Cold) | 5.414 (4.927,6.107) | - | residuals |
| | TempCat(Hot) | 3.295 (2.986,3.659) | - | residuals |
| **Repeatabilities** | Hot | 0.085 (0.045,0.131) | - | - |
| | Benign | 0.09 (0.053,0.151) | - | - |
| | Cold | 0.082 (0.047,0.133) | - | - |
| **Heritabilities** | Hot | 0.035 (0.018,0.076) | - | - |
| | Benign | 0.038 (0.017,0.081) | - | - |
| | Cold | 0.027 (0.016,0.071) | - | - |
| **Evolvabilities** | Hot | 0.017 (0.008,0.028) | - | - |
| | Benign | 0.027 (0.012,0.046) | - | - |
| | Cold | 0.027 (0.017,0.068) | - | - |

Estimate: Posterior mean is used for fixed effects and posterior mode is used for random effects

**Table S9: Results from animal model of neck surface temperature (character-state).**

| Type | Term | Estimate (CI) | pMCMC | Level |
|---|---|---|---|---|
| **Fixed effects** | Intercept | 32.8 (31.85,33.78) | **0.001** | - |
| | Temp__zcat | 1.3 (1.02,1.6) | **0.001** | - |
| | TempCat(Cold) | -4.14 (-5.09,-3.33) | **0.001** | - |
| | TempCat(Hot) | 5.19 (4.48,5.89) | **0.001** | - |
| | poly(Daytime_z, 2)1 | 17.28 (0.15,34.27) | **0.043** | - |
| | poly(Daytime_z, 2)2 | -28.71 (-38.46,-19.1) | **0.001** | - |
| | SubSpe(ZB) | -0.51 (-1.14,0.15) | 0.126 | - |
| | SubSpe(KR) | 0.5 (-0.31,1.36) | 0.254 | - |
| | SubSpe(Hybrid) | -0.02 (-0.5,0.44) | 0.875 | - |
| | Temp__zcat:TempCat(Cold) | -0.28 (-0.85,0.15) | 0.278 | - |
| | Temp__zcat:TempCat(Hot) | 0.89 (0.39,1.33) | **0.001** | - |
| | TempCat(Cold):SubSpe(ZB) | 0.17 (-0.76,1.33) | 0.746 | - |
| | TempCat(Hot):SubSpe(ZB) | 0.24 (-0.74,1.25) | 0.654 | - |
| | TempCat(Cold):SubSpe(KR) | -0.44 (-1.76,0.78) | 0.483 | - |
| | TempCat(Hot):SubSpe(KR) | -1.43 (-2.6,-0.26) | **0.016** | - |
| | TempCat(Cold):SubSpe(Hybrid) | -0.15 (-0.95,0.56) | 0.742 | - |
| | TempCat(Hot):SubSpe(Hybrid) | -0.26 (-0.87,0.37) | 0.426 | - |
| **Random effects** | TempCat(Benign) | 0.329 (0.142,0.53) | - | ID |
| | TempCat(Cold) | 0.511 (0.219,1.088) | - | ID |
| | TempCat(Hot) | 0.248 (0.112,0.435) | - | ID |
| | TempCat(Benign) | 0.19 (0.13,0.496) | - | animal |
| | TempCat(Cold) | 0.364 (0.148,0.75) | - | animal |
| | TempCat(Hot) | 0.249 (0.137,0.532) | - | animal |
| | year | 0.346 (0,3.125) | - | year |
| | enclosure | 0.17 (0.085,0.32) | - | enclosure |
| | date | 0.917 (0.442,1.387) | - | date |
| | TempCat(Benign) | 4.636 (4.204,5.1) | - | residuals |
| | TempCat(Cold) | 7.485 (6.727,8.413) | - | residuals |
| | TempCat(Hot) | 3.953 (3.579,4.447) | - | residuals |
| **Repeatabilities** | Hot | 0.088 (0.043,0.128) | - | - |
| | Benign | 0.082 (0.045,0.122) | - | - |
| | Cold | 0.097 (0.048,0.141) | - | - |
| **Heritabilities** | Hot | 0.036 (0.017,0.079) | - | - |
| | Benign | 0.032 (0.016,0.069) | - | - |
| | Cold | 0.035 (0.013,0.072) | - | - |
| **Evolvabilities** | Hot | 0.017 (0.01,0.037) | - | - |
| | Benign | 0.018 (0.012,0.046) | - | - |
| | Cold | 0.044 (0.018,0.091) | - | - |

Estimate: Posterior mean is used for fixed effects and posterior mode is used for random effects

**Table S10: Results from animal model of head-neck differences in surface temperature (random regression).**

In this model the difference between head and neck temperature at each thermal image was used a the response variable.

| Type | Term | Estimate (CI) | pMCMC | Level |
|---|---|---|---|---|
| **Fixed effects** | Intercept | 0.7 (0.23,1.2) | **0.016** | - |
| | SubSpe(ZB) | -0.17 (-0.62,0.31) | 0.44 | - |
| | SubSpe(KR) | -0.69 (-1.31,-0.11) | **0.027** | - |
| | SubSpe(Hybrid) | -0.18 (-0.52,0.18) | 0.317 | - |
| | poly(Daytime_z, 2)1 | -6.04 (-14.15,1.09) | 0.138 | - |
| | poly(Daytime_z, 2)2 | 3.11 (-1.79,8.33) | 0.222 | - |
| | TempCon:TempDir(Dec) | 0.48 (-0.99,1.97) | 0.514 | - |
| | TempCon:TempDir(Inc) | -1.79 (-2.36,-1.21) | **0.001** | - |
| | TempCon:TempDir(Dec):SubSpe(ZB) | -1.17 (-3.38,0.7) | 0.262 | - |
| | TempCon:TempDir(Inc):SubSpe(ZB) | 0.39 (-0.61,1.44) | 0.475 | - |
| | TempCon:TempDir(Dec):SubSpe(KR) | 1.41 (-0.97,3.95) | 0.258 | - |
| | TempCon:TempDir(Inc):SubSpe(KR) | 1.18 (0,2.43) | 0.051 | - |
| | TempCon:TempDir(Dec):SubSpe(Hybrid) | 0.63 (-0.78,2.13) | 0.418 | - |
| | TempCon:TempDir(Inc):SubSpe(Hybrid) | 0.27 (-0.38,1) | 0.466 | - |
| **Random effects** | (Intercept) | 0.139 (0.082,0.246) | - | ID |
| | TempCon:TempDir(Dec) | 4.511 (1.321,9.367) | - | ID |
| | TempCon:TempDir(Inc) | 0.365 (0.177,0.981) | - | ID |
| | (Intercept) | 0.119 (0.075,0.232) | - | animal |
| | TempCon:TempDir(Dec) | 0.584 (0.154,3.543) | - | animal |
| | TempCon:TempDir(Inc) | 0.356 (0.143,0.84) | - | animal |
| | enclosure | 0.001 (0,0.022) | - | enclosure |
| | date | 0.049 (0.013,0.098) | - | date |
| | year | 0.133 (0.027,0.784) | - | year |
| | residuals | 2.134 (1.997,2.247) | - | residuals |
| **Correlations** | TempCon:TempDir(Dec):(Intercept).animal | -0.26 (-0.68,0.32) | 0.53 | - |
| | TempCon:TempDir(Inc):(Intercept).animal | -0.52 (-0.76,-0.11) | **0.021** | - |
| | TempCon:TempDir(Inc):TempCon:TempDir(Dec).animal | 0.39 (-0.47,0.77) | 0.659 | - |
| | TempCon:TempDir(Dec):(Intercept).ID | 0.14 (-0.42,0.54) | 0.666 | - |
| | TempCon:TempDir(Inc):(Intercept).ID | -0.53 (-0.81,-0.2) | **0.01** | - |
| | TempCon:TempDir(Inc):TempCon:TempDir(Dec).ID | -0.46 (-0.77,0.36) | 0.456 | - |
| **Repeatabilities** | Intercept | 0.103 (0.072,0.153) | - | - |
| **Heritabilities** | Intercept | 0.049 (0.028,0.087) | - | - |
| **Evolvabilities** | Intercept | 31.002 (19.536,60.616) | - | - |

Estimate: Posterior mean is used for fixed effects and posterior mode is used for random effects

#### Differences among subspecies in morphology and climatic origin (Supplementary table 11-14)

**Table S11: Varimax rotated loadings of the PCA on 19 climatic variables.**

| | Comp.1 | Comp.2 | Comp.3 | Comp.4 |
|---|---|---|---|---|
| **anmean** | 0.3958827 | 0.0463324 | -0.0452534 | 0.0108890 |
| **Tdiurnalrange** | -0.0194926 | -0.4359334 | -0.1171183 | 0.1543744 |
| **Isothermality** | -0.0773870 | 0.3691935 | 0.0071038 | 0.0368339 |
| **Tseasonality** | 0.0493430 | -0.3880711 | 0.0928855 | -0.1264733 |
| **Tmax** | 0.3788211 | -0.1887792 | -0.0100697 | 0.0437655 |
| **Tmin** | 0.3081019 | 0.3323155 | 0.0086369 | 0.0043158 |
| **Tannualrange** | 0.0556095 | -0.4771859 | -0.0170450 | 0.0353619 |
| **Twet** | 0.3194865 | -0.0857049 | 0.0172106 | -0.1373188 |
| **Tdry** | 0.3710498 | 0.1720965 | 0.0231495 | 0.0554371 |
| **Twarm** | 0.3848324 | -0.1222385 | 0.0058273 | -0.0194566 |
| **Tcold** | 0.3240783 | 0.2337892 | -0.0813078 | 0.0977301 |
| **Pannual** | -0.0422437 | 0.0252377 | 0.0832177 | 0.4510177 |
| **Pwetm** | -0.0274255 | 0.0351452 | -0.1192616 | 0.4947200 |
| **Pdrym** | 0.0118639 | 0.0298580 | 0.5715990 | 0.0064004 |
| **Pseasonality** | 0.0917643 | 0.0319897 | -0.4386426 | 0.0557821 |
| **Pwetq** | -0.0389370 | -0.0022855 | -0.0980940 | 0.5198045 |
| **Pdryq** | 0.0110190 | 0.0565000 | 0.5685889 | 0.0015389 |
| **Pwarm** | -0.2498253 | 0.1100255 | 0.0046405 | 0.1326729 |
| **Pcold** | 0.1578871 | -0.1465692 | 0.3072379 | 0.4295756 |

**Table S12: Results from model of the difference between head and neck surface temperature with increasing/decreasing temperatures (random regression)**

In this model the difference between head and neck temperature (head-neck) at each thermal image was used a the response variable.

| Type | Term | Estimate (CI) | pMCMC | Lev |
|---|---|---|---|---|
| **Fixed effects** | Intercept | 0.69 (0.25,1.19) | **0.013** | - |
| | SubSpe(ZB) | -0.13 (-0.52,0.27) | 0.501 | - |
| | SubSpe(KR) | -0.64 (-1.13,-0.16) | **0.01** | - |
| | SubSpe(Hybrid) | -0.17 (-0.48,0.11) | 0.229 | - |
| | poly(Daytime_z, 2)1 | -6.31 (-13.77,1.65) | 0.11 | - |
| | poly(Daytime_z, 2)2 | 3.44 (-1.56,8.6) | 0.17 | - |
| | TempCon:TempDir(Dec) | 0.43 (-0.83,1.78) | 0.496 | - |
| | TempCon:TempDir(Inc) | -1.81 (-2.35,-1.27) | **0.001** | - |
| | TempCon:TempDir(Dec):SubSpe(ZB) | -1.15 (-3.2,0.68) | 0.229 | - |
| | TempCon:TempDir(Inc):SubSpe(ZB) | 0.39 (-0.53,1.35) | 0.413 | - |
| | TempCon:TempDir(Dec):SubSpe(KR) | 1.33 (-1.1,3.7) | 0.27 | - |
| | TempCon:TempDir(Inc):SubSpe(KR) | 1.23 (0.34,2.27) | **0.016** | - |
| | TempCon:TempDir(Dec):SubSpe(Hybrid) | 0.58 (-0.95,1.98) | 0.437 | - |
| | TempCon:TempDir(Inc):SubSpe(Hybrid) | 0.3 (-0.42,0.9) | 0.37 | - |
| **Random effects** | (Intercept) | 0.179 (0.1,0.291) | - | ID |
| | TempCon:TempDir(Dec) | 6.626 (2.527,9.416) | - | ID |
| | TempCon:TempDir(Inc) | 0.563 (0.201,1.189) | - | ID |
| | enclosure | 0.001 (0,0.024) | - | enc |
| | date | 0.048 (0.014,0.1) | - | dat |
| | year | 0.101 (0.031,0.813) | - | yea |
| | residuals | 2.138 (2.024,2.284) | - | resi |
| **Fixed effect contrasts** | TempCon:TempDir(Dec):SubSpe(KR) vs TempCon:TempDir(Dec):SubSpe(ZB) | 1.99 (-0.36,5.26) | 0.093 | - |
| | TempCon:TempDir(Inc):SubSpe(KR) vs TempCon:TempDir(Inc):SubSpe(ZB) | 1.99 (-0.36,5.26) | 0.178 | - |
| **Correlations** | TempCon:TempDir(Dec):(Intercept).ID | 0.17 (-0.29,0.57) | 0.574 | - |
| | TempCon:TempDir(Inc):(Intercept).ID | -0.65 (-0.82,-0.28) | **0.002** | - |
| | TempCon:TempDir(Inc):TempCon:TempDir(Dec).ID | -0.18 (-0.79,0.25) | 0.4 | - |

Estimate: Posterior mean is used for fixed effects and posterior mode is used for random effects

**Table S13: Results from model of the change in head surface temperature with increasing/decreasing temperatures (random regression) while accounting for body mass.**

| Type | Term | Estimate (CI) | pMCMC | Level |
|---|---|---|---|---|
| Fixed effects | Intercept | 30.93 (29.84,31.81) | **0.001** | - |
| | SubSpe(ZB) | -0.32 (-0.96,0.41) | 0.374 | - |
| | SubSpe(KR) | 0.07 (-0.74,0.72) | 0.827 | - |
| | SubSpe(Hybrid) | -0.02 (-0.43,0.45) | 0.926 | - |
| | Mass_z | -0.15 (-0.4,0.07) | 0.216 | - |
| | poly(Daytime_z, 2)1 | 14.82 (1.76,29.42) | **0.043** | - |
| | poly(Daytime_z, 2)2 | -27.77 (-35.48,-19.36) | **0.001** | - |
| | TempCon:TempDir(Dec) | -5.79 (-8.11,-3.53) | **0.001** | - |
| | TempCon:TempDir(Inc) | 10.31 (9.01,11.51) | **0.001** | - |
| | SubSpe(ZB):Mass_z | 0.09 (-0.39,0.7) | 0.757 | - |
| | SubSpe(KR):Mass_z | 0.69 (-0.28,1.63) | 0.162 | - |
| | SubSpe(Hybrid):Mass_z | 0.41 (-0.03,0.85) | 0.075 | - |
| | Mass_z:poly(Daytime_z, 2)1 | -4.6 (-14.11,4.82) | 0.338 | - |
| | Mass_z:poly(Daytime_z, 2)2 | -0.54 (-6.31,5.43) | 0.83 | - |
| | TempCon:TempDir(Dec):SubSpe(ZB) | 0.26 (-2.69,3.91) | 0.898 | - |
| | TempCon:TempDir(Inc):SubSpe(ZB) | -0.93 (-2.76,0.71) | 0.28 | - |
| | TempCon:TempDir(Dec):SubSpe(KR) | -0.66 (-4.32,2.33) | 0.69 | - |
| | TempCon:TempDir(Inc):SubSpe(KR) | -1.32 (-2.71,0.11) | 0.067 | - |
| | TempCon:TempDir(Dec):SubSpe(Hybrid) | -0.03 (-2.27,1.98) | 0.973 | - |
| | TempCon:TempDir(Inc):SubSpe(Hybrid) | -0.53 (-1.49,0.47) | 0.288 | - |
| | TempCon:TempDir(Dec):Mass_z | -0.46 (-1.85,1.04) | 0.539 | - |
| | TempCon:TempDir(Inc):Mass_z | 0.31 (-0.14,0.77) | 0.182 | - |
| | TempCon:TempDir(Dec):SubSpe(ZB):Mass_z | -1.64 (-4.22,0.8) | 0.206 | - |
| | TempCon:TempDir(Inc):SubSpe(ZB):Mass_z | 0.6 (-0.71,1.97) | 0.39 | - |
| | TempCon:TempDir(Dec):SubSpe(KR):Mass_z | -1.39 (-5.57,3.14) | 0.538 | - |
| | TempCon:TempDir(Inc):SubSpe(KR):Mass_z | -1.2 (-3.34,1.09) | 0.261 | - |
| | TempCon:TempDir(Dec):SubSpe(Hybrid):Mass_z | -0.89 (-2.66,1.2) | 0.363 | - |
| | TempCon:TempDir(Inc):SubSpe(Hybrid):Mass_z | -0.56 (-1.45,0.37) | 0.229 | - |
| Random effects | (Intercept) | 0.365 (0.184,0.685) | - | ID |
| | TempCon:TempDir(Dec) | 6.618 (1.456,13.752) | - | ID |
| | TempCon:TempDir(Inc) | 0.514 (0.205,1.601) | - | ID |
| | enclosure | 0.085 (0.001,0.169) | - | enclosure |
| | date | 0.617 (0.362,1.132) | - | date |
| | year | 0.462 (0.005,3.066) | - | year |
| | residuals | 4.173 (3.946,4.454) | - | residuals |

Estimate: Posterior mean is used for fixed effects and posterior mode is used for random effects

**Table S14: Results from model of the change in neck surface temperature with increasing/decreasing temperatures (random regression) while accounting for body mass.**

| Type | Term | Estimate (CI) | pMCMC | Level |
|---|---|---|---|---|
| Fixed effects | Intercept | 30.33 (29.45,31.29) | **0.001** | - |
| | SubSpe(ZB) | -0.13 (-0.87,0.64) | 0.715 | - |
| | SubSpe(KR) | 0.67 (-0.13,1.49) | 0.106 | - |
| | SubSpe(Hybrid) | 0.1 (-0.35,0.61) | 0.72 | - |
| | Mass_z | -0.16 (-0.42,0.11) | 0.213 | - |
| | poly(Daytime_z, 2)1 | 21.97 (5.86,37) | **0.008** | - |
| | poly(Daytime_z, 2)2 | -32.04 (-40.65,-22.37) | **0.001** | - |
| | TempCon:TempDir(Dec) | -6.29 (-8.73,-3.63) | **0.001** | - |
| | TempCon:TempDir(Inc) | 11.95 (10.54,13.31) | **0.001** | - |
| | SubSpe(ZB):Mass_z | -0.06 (-0.66,0.54) | 0.85 | - |
| | SubSpe(KR):Mass_z | 0.85 (-0.32,1.99) | 0.163 | - |
| | SubSpe(Hybrid):Mass_z | 0.47 (0,0.98) | 0.061 | - |
| | Mass_z:poly(Daytime_z, 2)1 | -1.05 (-11.76,9.47) | 0.806 | - |
| | Mass_z:poly(Daytime_z, 2)2 | 0.36 (-6.03,7.29) | 0.912 | - |
| | TempCon:TempDir(Dec):SubSpe(ZB) | 2.04 (-1.75,6.4) | 0.328 | - |
| | TempCon:TempDir(Inc):SubSpe(ZB) | -0.94 (-2.68,0.94) | 0.338 | - |
| | TempCon:TempDir(Dec):SubSpe(KR) | -2.23 (-6.38,1.66) | 0.28 | - |
| | TempCon:TempDir(Inc):SubSpe(KR) | -2.36 (-3.86,-0.72) | **0.002** | - |
| | TempCon:TempDir(Dec):SubSpe(Hybrid) | -0.78 (-3.19,1.69) | 0.554 | - |
| | TempCon:TempDir(Inc):SubSpe(Hybrid) | -0.46 (-1.51,0.71) | 0.411 | - |
| | TempCon:TempDir(Dec):Mass_z | 0.08 (-1.52,1.84) | 0.923 | - |
| | TempCon:TempDir(Inc):Mass_z | 0.2 (-0.37,0.7) | 0.475 | - |
| | TempCon:TempDir(Dec):SubSpe(ZB):Mass_z | -2.31 (-5.28,0.7) | 0.114 | - |
| | TempCon:TempDir(Inc):SubSpe(ZB):Mass_z | 0.7 (-0.89,2.05) | 0.352 | - |
| | TempCon:TempDir(Dec):SubSpe(KR):Mass_z | -1.64 (-7.3,3.16) | 0.538 | - |
| | TempCon:TempDir(Inc):SubSpe(KR):Mass_z | -1.5 (-3.82,1.09) | 0.235 | - |
| | TempCon:TempDir(Dec):SubSpe(Hybrid):Mass_z | -1 (-3.23,1.17) | 0.363 | - |
| | TempCon:TempDir(Inc):SubSpe(Hybrid):Mass_z | -0.79 (-1.79,0.2) | 0.15 | - |
| Random effects | (Intercept) | 0.453 (0.221,0.819) | - | ID |
| | TempCon:TempDir(Dec) | 15.078 (5.394,24.954) | - | ID |
| | TempCon:TempDir(Inc) | 0.431 (0.164,1.545) | - | ID |
| | enclosure | 0.141 (0.045,0.292) | - | enclosure |
| | date | 0.668 (0.43,1.358) | - | date |
| | year | 0.009 (0.001,2.877) | - | year |
| | residuals | 5.177 (4.904,5.56) | - | residuals |

Estimate: Posterior mean is used for fixed effects and posterior mode is used for random effects

#### Evolutionary parameters of the head-neck difference in surface temperature (Supplementary tables 15-16)

##### Table S15: Evolutionary parameters of the head-neck difference in surface temperature (character-state)

These estimates are also available in **Table 8**.

| | Benign | Cold | Hot |
|---|---|---|---|
| **Additive genetic variance** | 0.151 (0.081,0.236) | 0.226 (0.115,0.512) | 0.186 (0.089,0.268) |
| **Repeatabilities** | 0.117 (0.082,0.16) | 0.173 (0.125,0.264) | 0.167 (0.102,0.208) |
| **Heritabilities** | 0.06 (0.03,0.09) | 0.062 (0.03,0.128) | 0.08 (0.041,0.122) |
| **Evolvabilities** | 247.12 (133.592,386.818) | 26.965 (13.791,61.17) | 150.675 (72.002,217.199) |

##### Table S16: Evolutionary parameters of the head-neck difference in surface temperature (random regression)

These estimates are also available in **Table 9**.

| | Intercept |
|---|---|
| **Additive genetic variance** | 0.119 (0.075,0.232) |
| **Repeatabilities** | 0.103 (0.072,0.153) |
| **Heritabilities** | 0.049 (0.028,0.087) |
| **Evolvabilities** | 31.002 (19.536,60.616) |

#### Supplementary Figures

**Figure S1: No evidence for active regulation of neck surface temperature.**

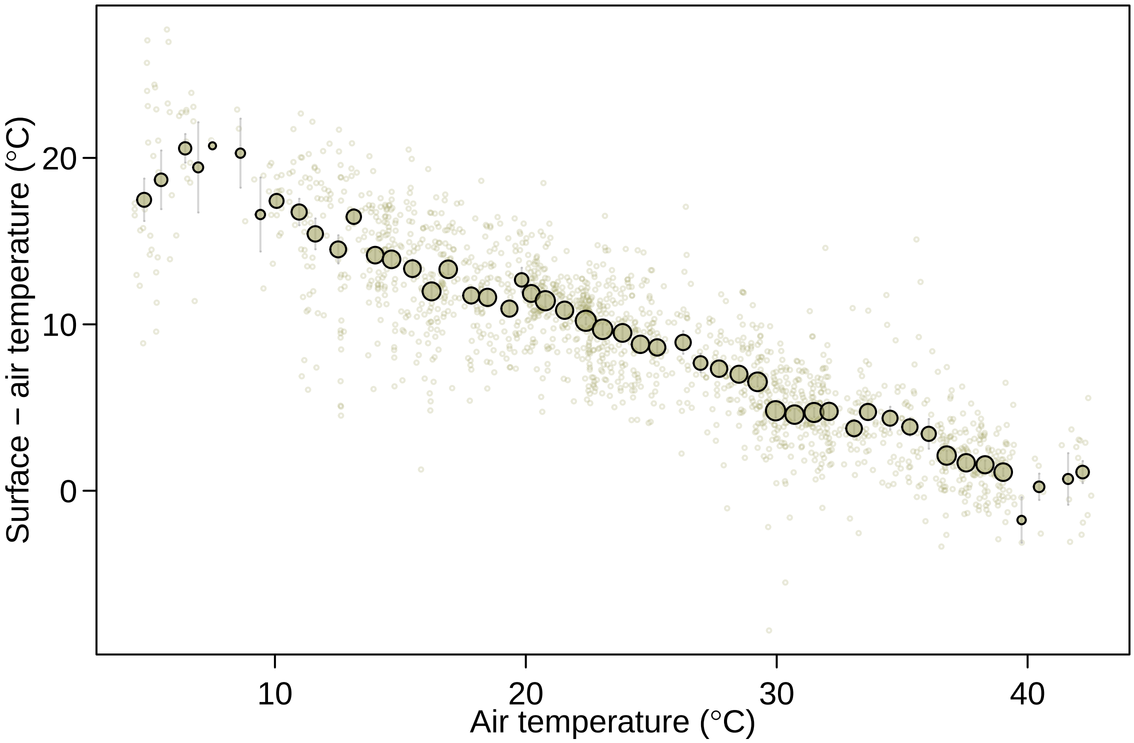

Larger points are averages with standard errors binned according to temperature. Point size illustrates relative number of individuals: smallest point = 1 and largest point = 65.

**Figure S2: Relationship between body mass and neck temperature across subspecies.**

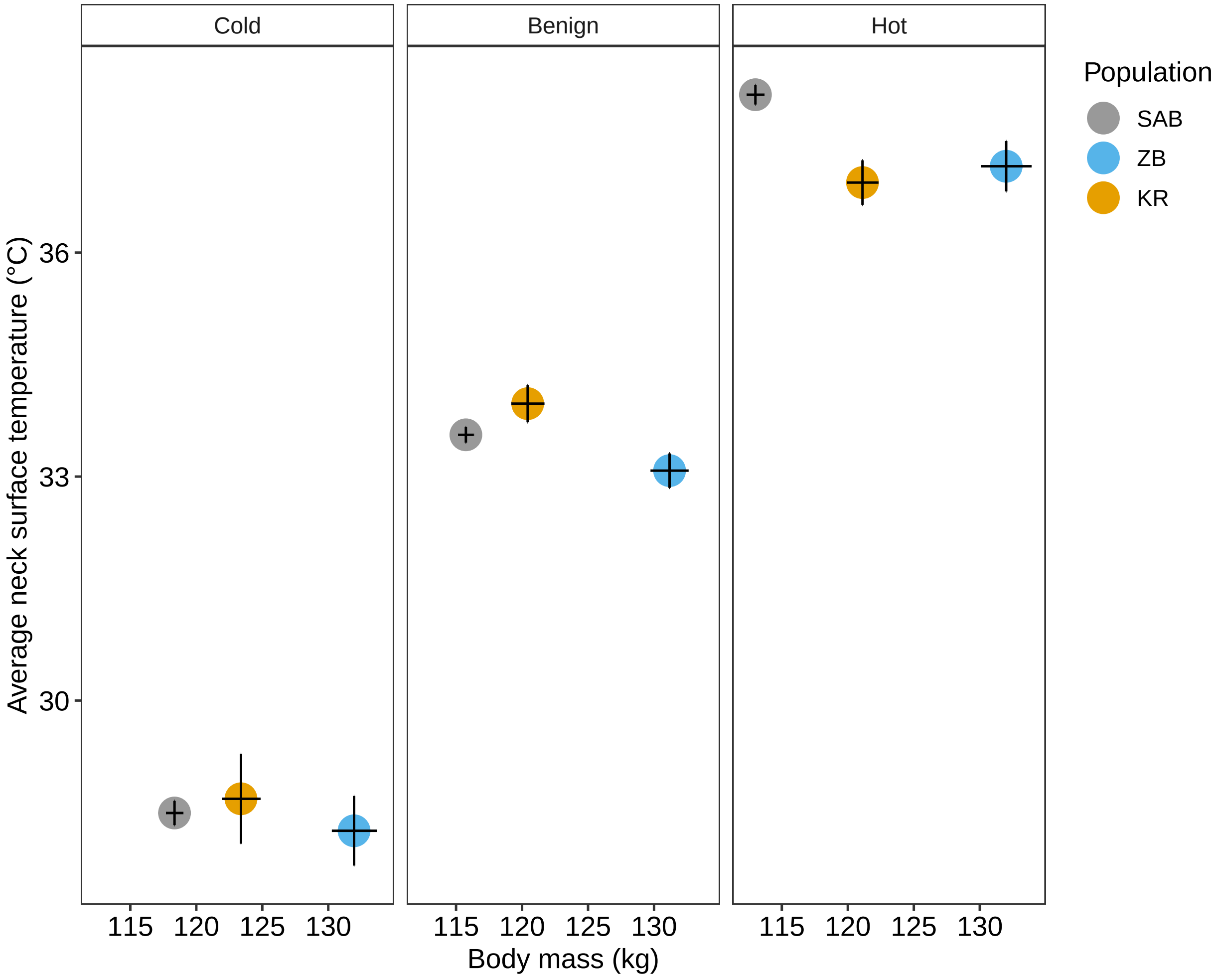

Each point is the mean standard errors as error bars.

**Figure S3: Relationship between body mass and neck temperature within sub-species.**

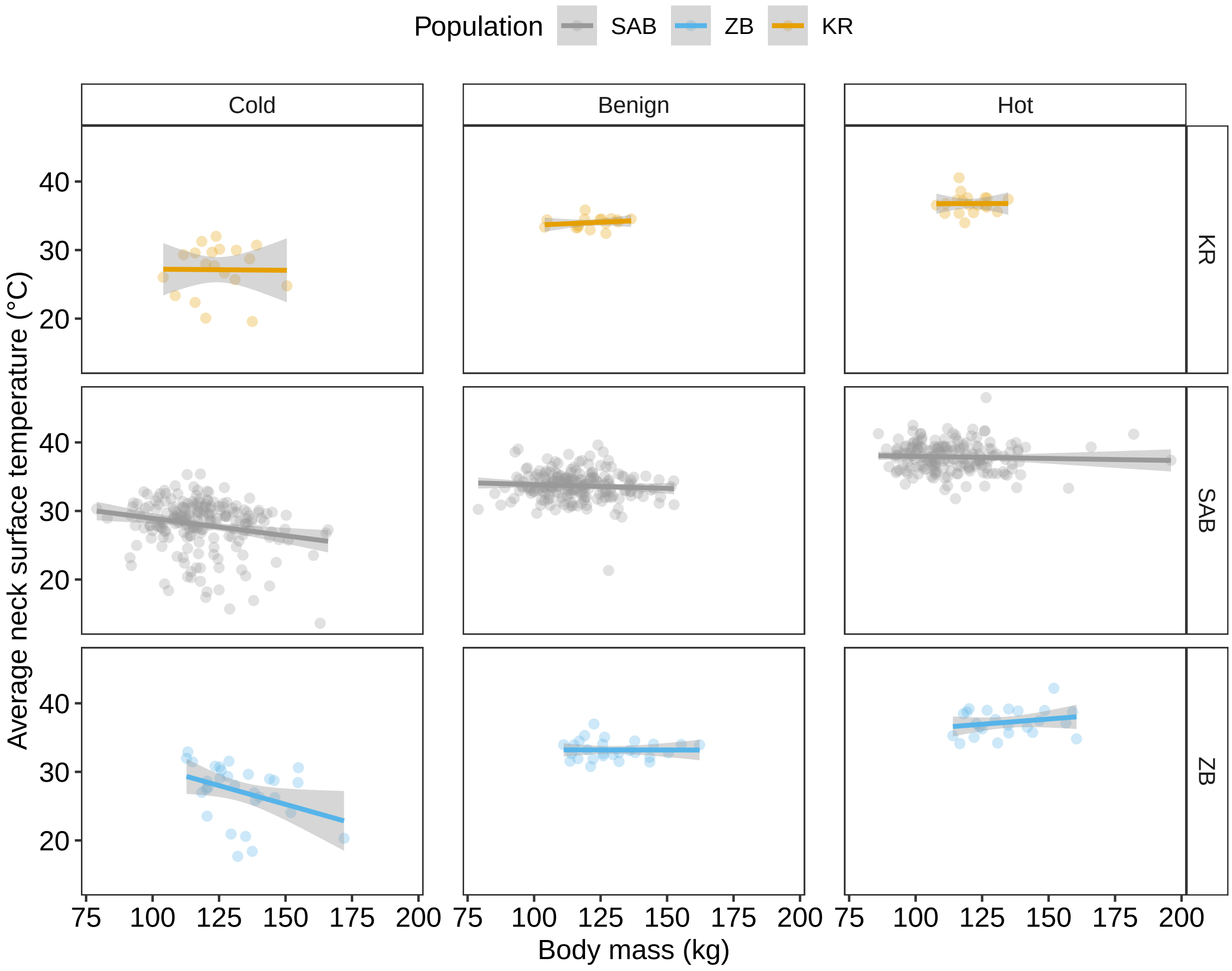

Each point is the individual mean.

#### Figure S4: Effect of air temperature on surface temperatures

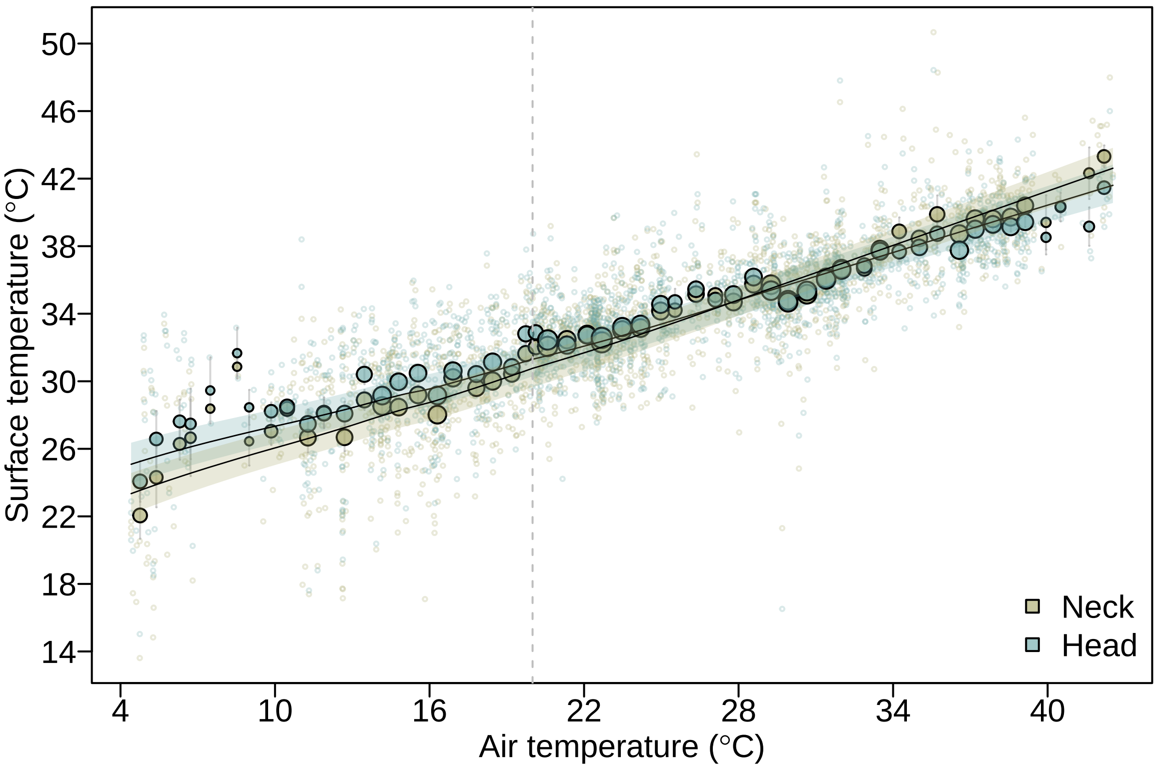

The featherless and vascularized body parts of the head and neck surface temperatures were sensitive to increases (nimages = 1895) and decreases (nimages = 849) from 20°C air temperatures (vertical line). The larger points are averages with standard errors binned according to temperature. Point size illustrates relative number of individuals: smallest point = 2 and largest point = 71. Fitted lines and 95% credible intervals (shaded area) were extracted from the statistical models (**Tables S1-S2**). See methods for further details.

**Figure S5: Four first principal components from PCA on bioclimatic variables in the ostrich range**

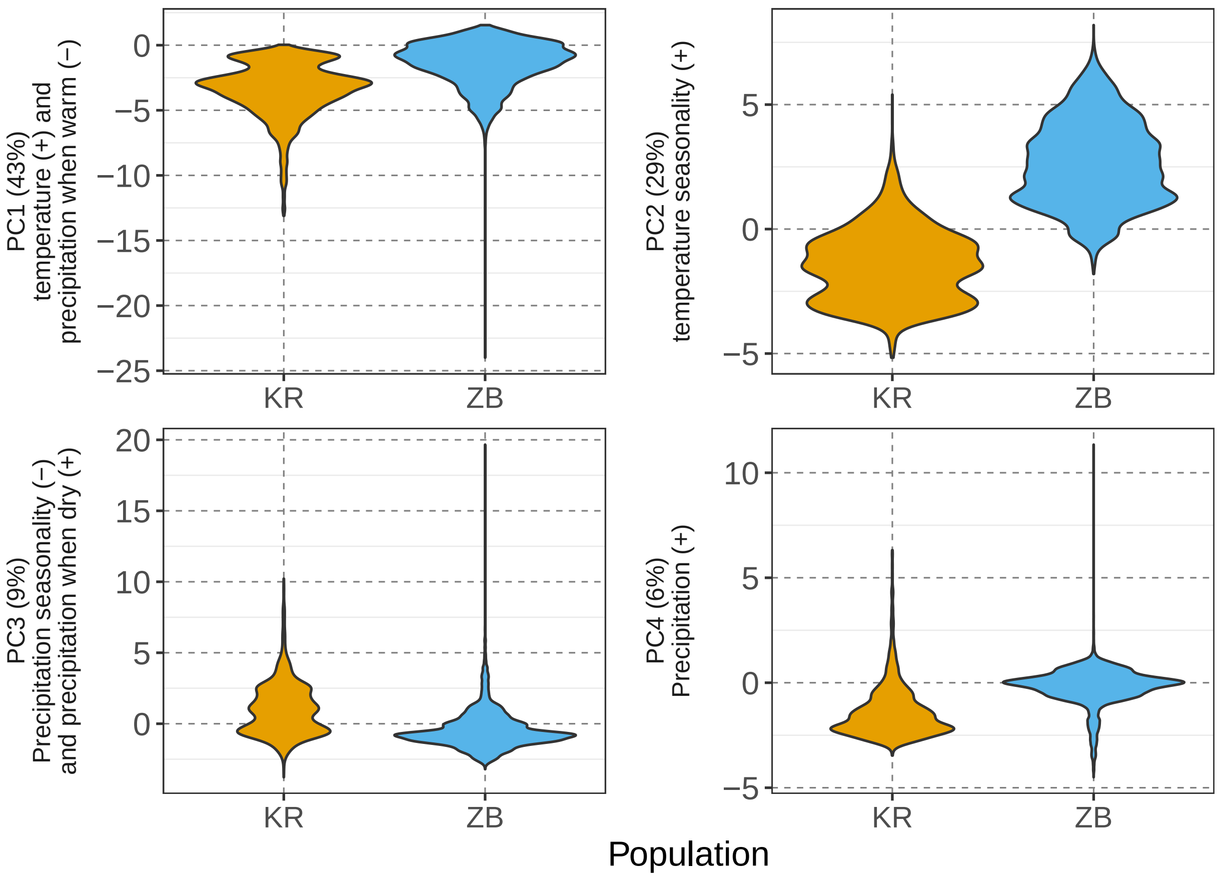

**Figure S6: Change in relative head temperature with increasing air temperature across subspecies**

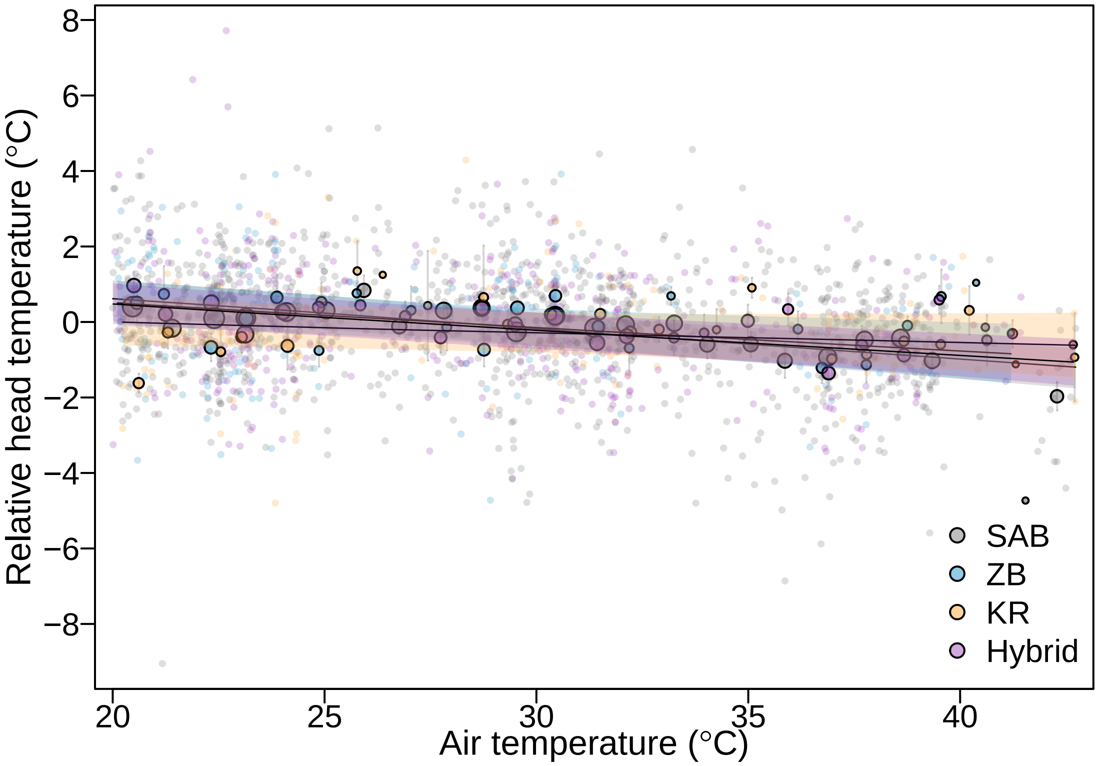

Fitted lines and 95% credible intervals (shaded area) of the change in relative head temperature with increasing air temperature. SAB, ZB and Hybrids individuals have very similar reduction in relative head temperature when air temperatures increase, while KR individuals show no reduction (**Table S11**). Larger points are averages with standard errors binned according to the temperature variable.

**Figure S7: Change in relative head temperature with decreasing air temperature across subspecies**

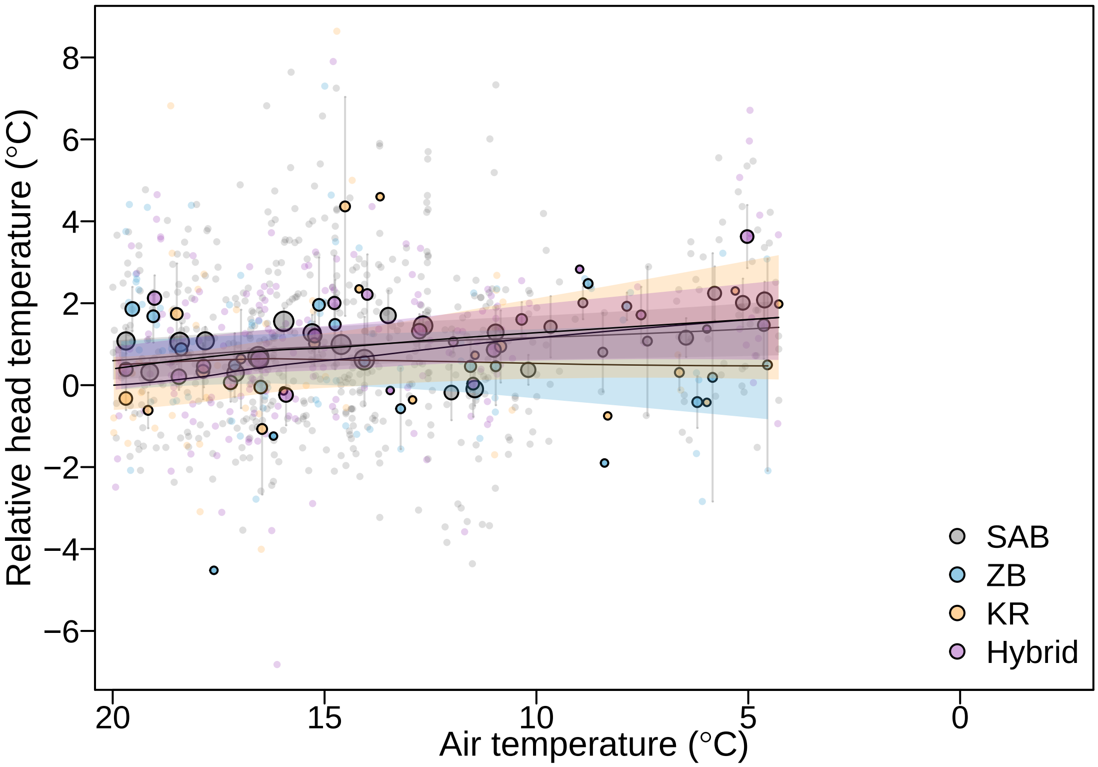

Fitted lines and 95% credible intervals (shaded area) of the change in relative head temperature with decreasing air temperature. Larger points are averages with standard errors binned according to the temperature variable.

**Figure S8: Change in feather, head and neck temepratures with increasing air temperature**

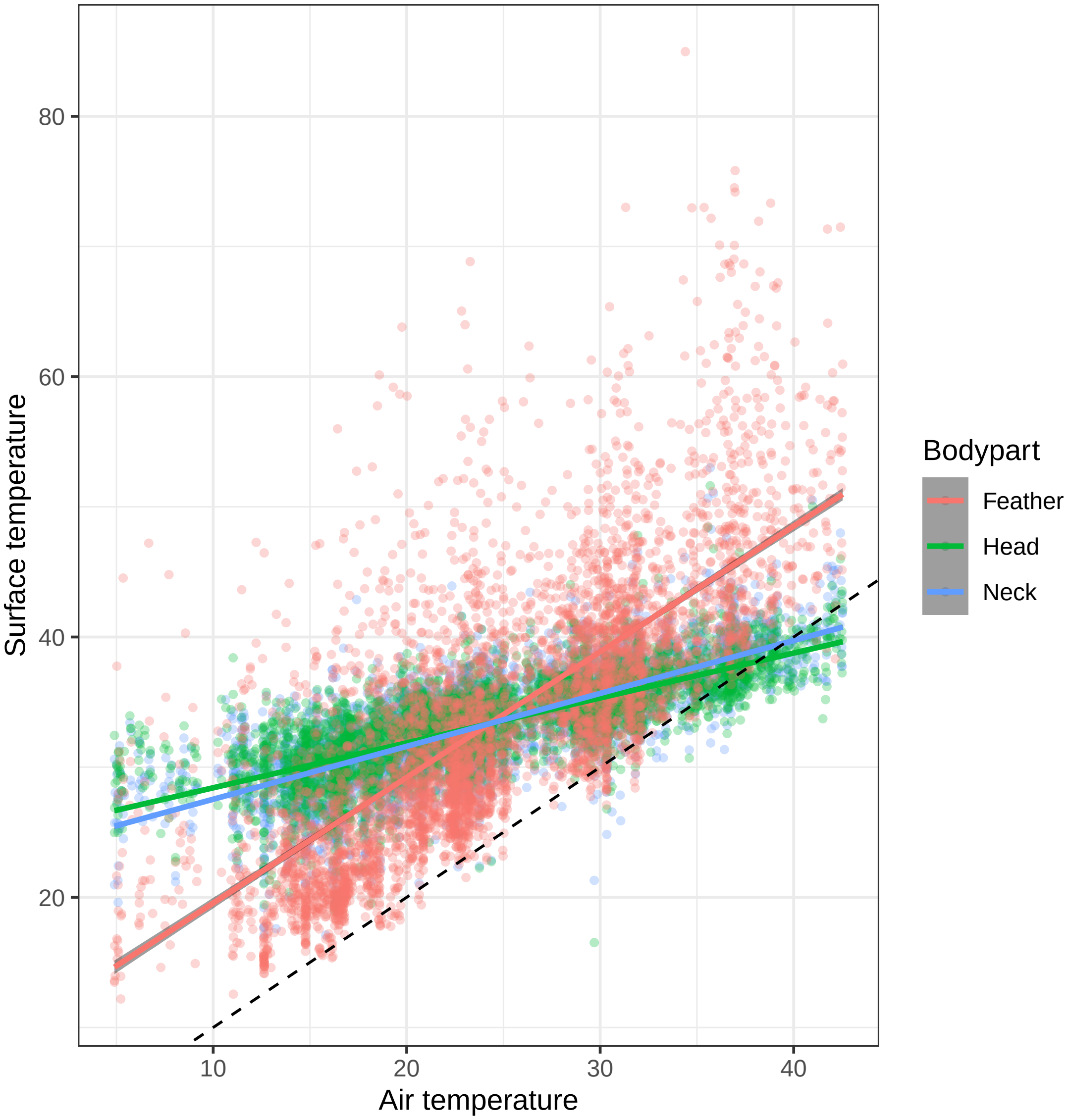

This comparison of the change in feather, head and neck temperatures with air temperature is based on 2451 pictures where all there measures where available. Feather temperature appear correlated with air temperature but highly variable. We believe this is because when animals are free ranging you have little control over events leading up to the image being taken (e.g. feathers could have been facing the sun or away from the sun). The punctuated black line is y = x.

**Figure S9: Impact of distance to ostrich during thermal imaging**

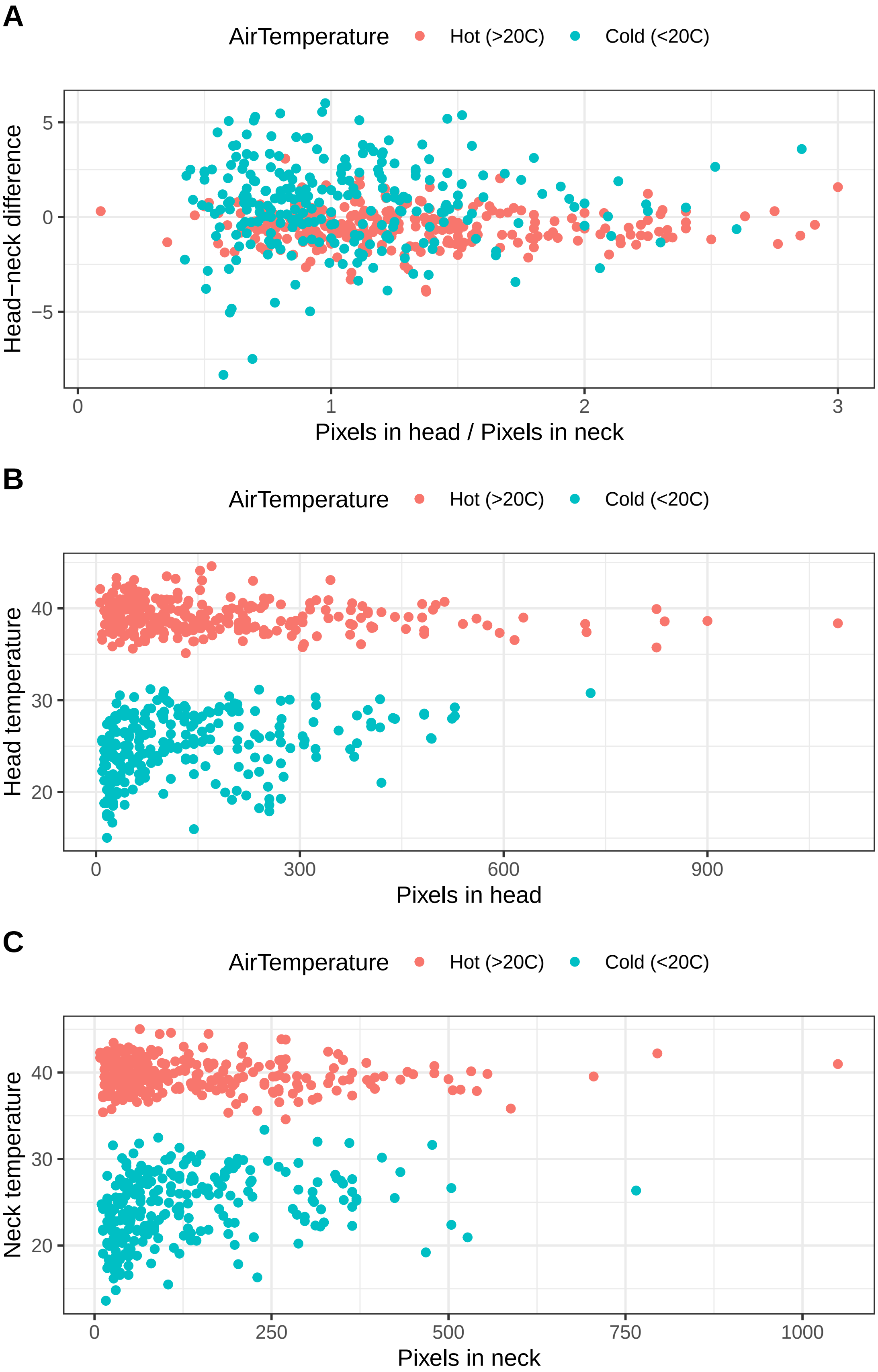

We used the number of pixels used to estimate surface temperatures of the head and neck as a proxy for distance in 534 pictures. Extracting this data from all of our images is a very time demanding task that was not possible to achieve. We extracted pixel information from ~25% of the pictures to investigate potential biases on measures of head-neck differences induced by the distance to birds. Using this subset of data, we found that our metric of head-neck surface temperature (A) is actually independent of the difference in number of pixels between head and neck. In pictures taken far away, both head and neck are small (B-C), resulting in any biases being accounted for by the head-neck metric. In contrast, if we just examined head or neck measures, we find more error in images where birds are further away. Taken together this suggests that the potential distance biases do not have a severe impact on our measures of head-neck temperature differences.

**Figure S10: Impact of angle during thermal imaging**

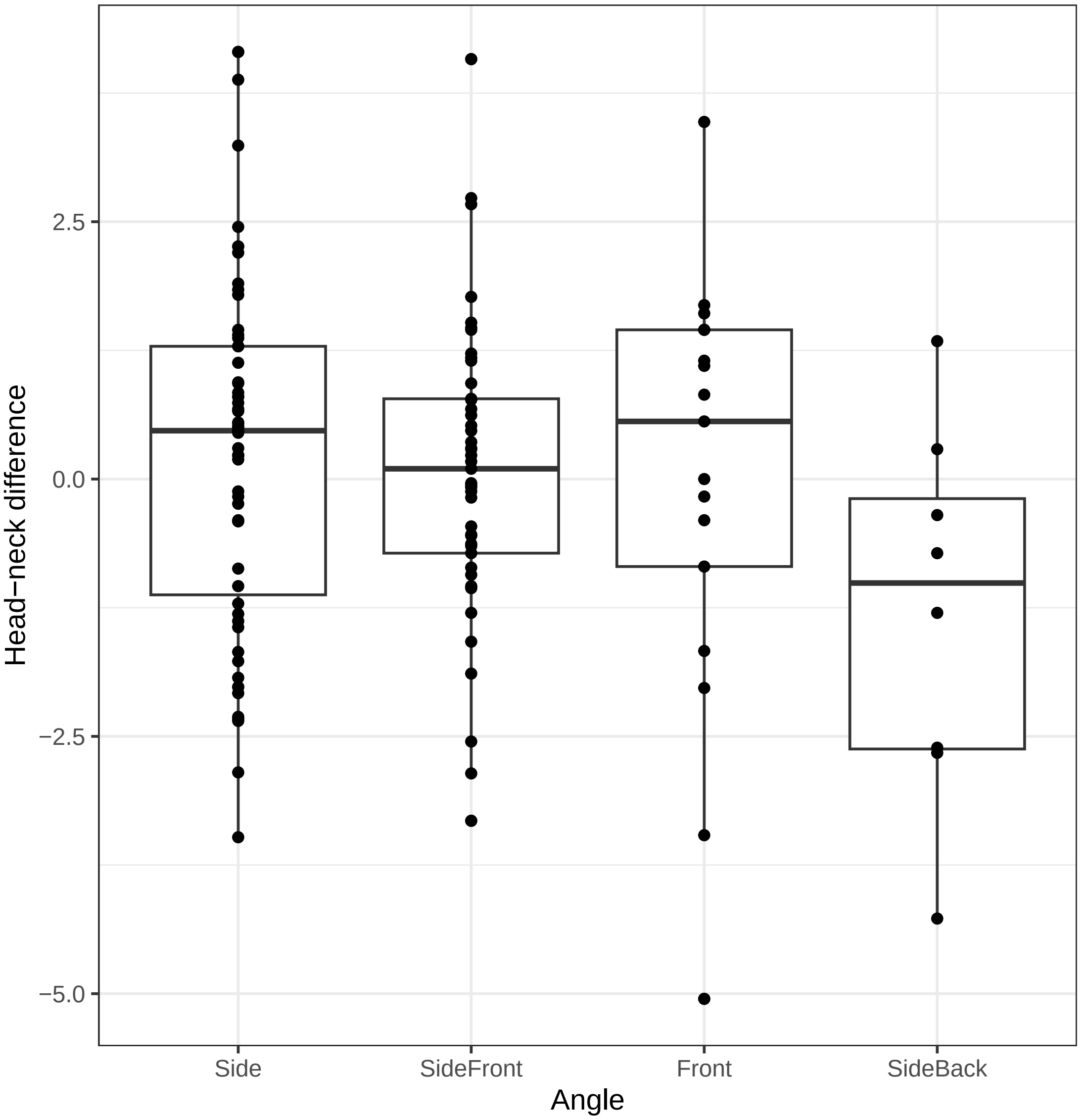

In ostriches the head and the neck move together synchronously. Ostriches also typically orientate themselves side on to people so that the side of the head and neck consistently face the investigator (Figure 1B). To quantify this, we recorded the angle of the head in a subset of pictures (n=130) and found 87% included the side of the head, and thereby the eye. In the remaining pictures, the front of the head faced the investigator, thereby including a bit of each eye and the bill. No pictures were taken of the back of head. We found no effect of the angle of the head on the head-neck difference ($F = 2.08$ $df = 3{,}126$, $P = 0.101$).
